# Molecular Cloning, *In Silico* Analysis and Expression of plasma membrane-associated NAR2 Protein, SaNAR2.2, from Euhalophyte *Suaeda altissima*

**DOI:** 10.1101/2025.03.09.642219

**Authors:** D.E. Khramov, A.O. Konoshenkova, O.I. Nedelyaeva, E.I. Rostovtseva, A.V. Ryabova, V.S. Volkov, L.G. Popova

**Author notes:** Correspondence (P.L.G.); (V.V.S.). (K.D.E.); (K.A.O.); (N.O.I.); (R.E.I.); (V.S.V.); (P.L.G.).

## Abstract

Plasma membrane-associated proteins of NAR2 family are involved in the uptake of nitrate by plant roots as partners of NRT2 family proteins, which carry out high-affinity nitrate transport across membranes of plant cells. In this study, the coding sequence of *NAR2* family gene, *SaNAR2.2,* was cloned from the euhalophyte *Suaeda altissima*. This species is able to withstand high concentrations of NaCl, up to 750 mM, in the nutrient medium. The protein SaNAR2.2, of 196 amino acids in size, with a calculated molecular weight of 21.7 kDa, has been characterized in terms of structure and phylogeny. According to the prediction of cellular localization and protein topology, SaNAR2.2 is a resident of plasma membrane. It has one transmembrane segment at the C-terminal part and a small C-end located in cytoplasm. The large N-terminal part of SaNAR2.2 protein is located in the extracellular space. On the phylogenetic tree of NAR2 family proteins, the protein from *S. altissima* lies in a clade with the proteins of NAR2.2 subfamily from other plants of Amaranthaceae family. We detected the interaction between SaNAR2.2 and putative partner proteins SaNRT2.1 and SaNRT2.5, which are the members of high-affinity nitrate transporter family NRT2 from *S. altissima*, using the bimolecular fluorescence complementation (BiFC) assay in *Nicotiana benthamiana* leaves. The expression of *SaNAR2.2* gene in organs of *S. altissima* plants grown at low (0.5 mM) or high (15 mM) nitrate concentrations in the nutrient medium, without NaCl or in the presence of increasing salt concentrations, was analyzed using RT-qPCR method. It has been found that *SaNAR2.2* is expressed nearly exclusively in roots of *S. altissima*. As the concentrations of NaCl in the nutrient medium increased (up to 500 mM), the expression of *SaNAR2.2* decreased at high external nitrate concentrations and significantly increased, up to ten-fold, at low nitrate concentrations; this points to the significant role of SaNAR2.2 protein under salinity conditions as a participant of high-affinity nitrate transport system. SaNAR2.2 is the first components of a high affinity nitrate transport system characterized in a terrestrial halophytic plant.

## Introduction

Nitrogen is one of the most important biogenic elements. Nitrogen is available to plants in various forms, including nitrate (NO_3_^̶^), ammonium (NH_4_^+^), and organic forms (amino acids) [1]; however, plants most often obtain nitrogen from the soil in the form of nitrate, which is the dominant form among nitrogen compounds in aerobic soils [2, 3]. Plants absorb nitrate from the soil solution using specialized transport mechanisms that function in the plasma membranes of root cells. Currently, four families of nitrate transport proteins are known - NRT1/PTR/NPF (Nitrate Transporter 1/Peptide Transporter Family/Nitrate Peptide Family), NRT2 (Nitrate Transporter 2), CLC (Chloride Channel Family), SLAC/SLAH (SLow Anion Channel/SLow Anion channel Homolog) [4–7], but only the proteins of NPF and NRT2 families identified as significantly involved in nitrate uptake by plant roots [8].

The availability of nitrate in the soil varies significantly, and this has led to the development of transport systems in plants with different affinities for nitrate [1]. Low-affinity transport systems belonging to NRT1/PTR/NPF family make a major contribution to nitrate uptake at high external nitrate concentrations (above 1 mM) [1, 9]. At low nitrate concentrations in the external environment (from 1 µM to 0.5 mM), high-affinity nitrate transporters of NRT2 family play a decisive role in plant nitrogen nutrition [10–13]. Nitrate transporters of NRT2 family have been identified in a wide range of plants including *Arabidopsis thaliana* [14], barley (*Hordeum vulgare*) [15], wild soybean (*Glycine soja*) [16], rice (*Oryza sativa*) [17], spinach (*Spinacea oleracea*) [18], and others. The NRT2 family proteins have been most extensively studied in the model plant *Arabidopsis thaliana* [19–21].

However, the essential feature of NRT2 proteins is their capability to transfer NO_3_^−^ across the membrane when provided a complex formed with a partner protein from NAR2/NRT3 family (Nitrate Assimilation Related Protein/ Nitrate Transporter 3) [22–26]. NAR2 is a family of small membrane-associated proteins that are important components of high-affinity nitrate uptake system in plants [27]. Genes belonging to NAR2 family have been identified in a number of plant species: three genes (*HvNAR2.1* – *HvNAR2.3*) have been found in barley [28], two genes were found in both *Arabidopsis* [24] and rice [29], and one NAR2 gene was revealed in chrysanthemum (*Chrysanthemum morifolium*) [30], maize (*Zea mays*) [31, 32], seagrass *Zostera marina* [33], wheat (*Triticum aestivum*) [34], and spinach [18].

Interaction between NRT2 and NAR2 proteins has been well documented. In *A. thaliana*, two members of the NAR2 family, AtNAR2.1 and AtNAR2.2, have been identified [24]. It has been shown that, with the exception of AtNRT2.7, all other members of the high-affinity NRT2 family of *A. thaliana* interact with AtNAR2.1 [26]. In experiments with *Arabidopsis*, obtained data indicated that the active transport form of the NRT2/NAR2 protein complex is likely a tetramer located in the plasma membrane; the complex presumably consists of two subunits, each of them was a heterodimer of NRT2.1 and NAR2.1 [25, 35]. The involvement of NAR2 proteins in nitrate transport as partners of NRT2 proteins was proved for other plant species, including barley [15], wild soybean (*Glycine soja*) [16], spinach [18], rice [27], rapeseed (*Brassica napus*) [36], seagrass *Zostera marina* [37].

In addition to its role as a component of high affinity transport systems, NAR2 protein has many other functions [28]. NAR2 proteins are necessary for the stabilization and localization of NRT2 transporters in the plasma membrane [38]. Co-localization of NRT2 and NAR2 proteins in the plasma membrane of root cells has been demonstrated for barley [39] and for *Arabidopsis* proteins [25, 26]. For the proteins from *Oryza sativa*, it has been shown that OsNAR2.1 targets OsNRT2.3 to the plasma membrane [40]. NAR2 proteins are also involved in suppressing lateral root formation in response to a high sucrose-to-nitrogen ratio in the environment [41], i.e., the two-component system NRT2/NAR2 is likely involved in the signaling pathway that regulates lateral root architecture [42, 43].

The available information on plant nitrate transporters has been obtained mainly from salt-sensitive glycophyte species, particularly from the model plant *A. thaliana*. Data about nitrate transporters in salt tolerant halophytic plants are very limited. Currently, we only know one example of the high affinity transport system NRT2/NAR2 discovered in an extremely salt-tolerant halophyte, namely, ZosmaNRT2 and ZosmaNAR2 from the seagrass *Zostera marina* [33, 37]. Soil salinization with sodium chloride reduces the availability of nitrate for plants. The main reason for this is the direct competition between the anions NO_3_^−^ and Cl^−^ for high-affinity nitrate transporters [44, 45]. Under these conditions, NO_3_^−^ uptake and nitrogen assimilation in glycophytes are suppressed [46, 47]. In highly salt-tolerant halophytes growing on saline substrates, nitrate-transporting plasma membrane proteins are able to function while under conditions in contact with an external solution with a high chloride concentration. This suggests the presence of structural features in halophyte proteins that allow them to bind nitrate and transport it across the membrane when the concentration of this anion in the surrounding solution is much lower than the concentration of chloride ions. It can also be assumed that under conditions of nitrate deficiency and simultaneous salinization of the nutrient medium with sodium chloride, high-affinity nitrate transporters play a key role in the absorption of nitrate ions by roots.

In our previous research, we cloned the coding regions of two genes from the euhalophyte *Suaeda altissima*, *SaNRT2.1* (OR909030.1) and *SaNRT2.5* (OR828748.1), which belong to the high-affinity nitrate transporter gene family *NRT2* [48]. Many representatives of the genus *Suaeda* (Amaranthaceae/Chenopodiaceae) have been identified as high salt tolerant species [49, 50]. *S. altissima* is one of the most salt tolerant plant species. This species is able to proliferate high concentrations of NaCl in the nutrient medium (up to 750 mM), when at NaCl concentrations up to 250 mM, the growth of *S. altissima* was not reduced, but stimulated [51]. Under these conditions, chloride competes with nitrate for binding sites of nitrate transporters, so the ability of halophytes, including *S. altissima*, to complete their life cycle in highly saline and simultaneously nitrate-poor soils may be due to the unique properties of the high-affinity nitrate transporters of these plants. The proteins SaNRT2.1 and SaNRT2.5 from *S. altissima* are orthologs of high-affinity nitrate transporters AtNRT2.1 and AtNRT2.5 from *A. thaliana*, respectively, which were functionally active only in complex with the protein AtNAR2.1 also known as AtNRT3.1 [26]. It is assumed that SaNRT2.1 and SaNRT2.5 also form functional complexes with proteins of NAR2/NRT3 family. The present work is aimed at identification of the coding sequence of the *NAR2* family gene in *S. altissima*, *SaNAR2.2*. The expression of this gene in *Suaeda* organs was examined at various concentrations of nitrate and chloride in the nutrient medium. Using the bimolecular fluorescence complementation (BiFC) method [52, 53], we demonstrated that protein SaNAR2.2 interacts with proteins SaNRT2.1 and SaNRT2.5 that belong to high affinity nitrate transporter family NRT2. To analyse the function of SaNAR2.2 protein and NRT2 family transporters from *S. altissima*, the coding sequences of *SaNAR2.2* and *SaNRT2.1/SaNRT2.5* were co-expressed in *Δynt1* mutant strain of *Hansenula (Ogatae) polymorpha* yeast. Methylotrophic yeast cells of *H. polymorpha* are capable of using nitrate as the only nitrogen source and possess the genes required for its uptake from the medium and assimilation [54]. The high-affinity nitrate uptake in *H. polymorpha* is mediated by a single transporter encoded by the *YNT1* gene (Yeast Nitrate Transporter 1) [55]. The knockout mutant *H. polymorpha* in this gene, *Δynt1*, is used to study heterologously expressed nitrate transporters from plants [37, 54, 56, 57].

## Materials and Methods

### 2.1. Plant Material

The euhalophyte *Suaeda altissima* (L.) Pall. was the object of the study for this research. *S. altissima* seeds collected from plants growing in the coastal zone of the salt lake Elton (Volgograd Region, Russia), were germinated in wet sand. The seedlings aged 14 days were transferred to 3-liter vessels with liquid nutrient medium (four plants per a vessel). The plants were grown in a hydroponic culture, on the aerated Robinson-Downton nutrient medium [58]. The plants were grown at two concentrations of NO_3_^−^ in the medium, low (0.5 mM) and high (15 mM), and four concentrations of Cl^−^ (0, 250, 500, and 750 mM) for each of the specified nitrate concentrations. Nitrate and chloride were added to the nutrient solution in the form of KNO_3_ and NaCl, respectively. Salt NaCl was added to the nutrient solution starting from the seventh day after the transfer of seedlings to the hydroponic culture. The concentration of NaCl in the nutrient solution gradually increased by 50-100 mM per day until the final concentrations of 250, 500, and 750 mM were reached. The control plants did not include NaCl in the nutrient medium. Seed germination and plant cultivation were carried out under 24°C and relative humidity of 60-70%. The plants were illuminated with high-pressure sodium lamps DNaZ_400 (Reflux, Novocherkassk, Russia) (light flux 300 µmol photons m^−2^ s^−1^, 16 hours light/8 hours dark).

The plants of *Nicotiana benthamiana* were grown from seeds in soil substrate at 23 ± 2 °C, relative air humidity of 70 ± 5%, photoperiod of 12 h per day, and light intensity of 120 µmol quanta/(m^2^·s) emitted from Phillips TL-D-58W/33-640 fluorescent lamps (Poland). For transient expression in *N. benthamiana*, forty-day-old plants were used.

### 2.2. Isolation of total RNA from plant material

To obtain total RNA from *S. altissima* organs, 6-week-old plants were used. Samples of plant material (roots, leaves, stems) with weight approximately 0.5 g were frozen in liquid nitrogen and stored at -80°C for the further RNA extraction. Total RNA from *S. altissima* organs was extracted by the hot phenol method [59] and treated with DNase I (Thermo Fisher Scientific, USA) to remove genomic DNA contaminants. The concentration and purity of the RNA preparations were determined spectrophotometrically by the A260/A280 ratio using the NanoDrop ND1000 device (Thermo Fisher Scientific, Inc., Waltham, MA, USA). One µg of total RNA in a total volume of 20µl was used as a template for the synthesis of the first strand of cDNA.

### 2.3. Primer design

For qRT-PCR experiments, primers were designed using primer Blast software, version 4.1.0 (https://www.ncbi.nlm.nih.gov/tools/primer-blast/). In other cases, SnapGene Viewer v.7.2. software was used for the primer design. All primers used are listed in Table S1 (Supporting material).

### 2.4. Yeast Strain and Vectors Used in This Study

Methylotrophic yeast *Hansenula polymorpha* double-auxotrophic strains DL-1 (*leu2 ura3* genotype) (wild-type strain, WT strain) and the mutant strain *Δynt1* (Δ*ynt1: BleoR/ZeoR, leu2, ura3*) with a deleted gene *YNT1* encoding the only high-affinity nitrate transporter in *H. polymorpha* were used in this study. The mutant strain was obtained by us earlier [57]. The WT strain and the mutant strain *Δynt1* were transformed with yeast integrative plasmids pCCUR2 and pCHLX [60] carrying the *URA3* and *LEU2* genes, respectively, to ensure the growth of the yeast strains in the selective media without additional nitrogen sources in the selective media, namely leucine and uracil, when performing complementation tests. Plasmid pCCUR2 was constructed by Michael Agaphonov (Moscow) by deletion of KpnI-XbaI fragment in the pCCUR1 plasmid [61]. Plasmids pCCUR2 and pCHLX were kindly provided by Michael Agafonov (Federal Research Center “Fundamentals of Biotechnology”, Russian Academy of Sciences, Moscow, Russia).

### 2.5. Cultivation of H. polymorpha WT Strain and Δynt1 Transformants

Cells of *H. polymorpha* WT strain and the mutant *Δynt1* strain were routinely grown in a rich YPD medium (1% yeast extract, 2% peptone, 2% glucose). After transformation with plasmid vector constructs, yeast cells were grown in a minimal synthetic SD medium (0.17% yeast nitrogen base without amino acids and ammonium sulfate, 2% glucose) with the addition of 0.5% (NH_4_)2SO4 as a nitrogen source. All agarized media contained 2% agar. Yeast transformants containing genomic inserts were verified using PCR.

### 2.6. Cloning of the full-length coding sequence SaNAR2.2

To clone the full-length coding sequence of the *NAR2* gene from *S. altissima*, the sequences of its 5’- and 3’-end fragments were initially determined using the Step-Out RACE method (kit No. SKS03, Evrogen, Moscow). The amplification of the 5’- and 3’-end fragments of the targeted *NAR2* transcript, as well as the amplification of the full-length coding sequence of *NAR2* from *S. altissima*, was carried out on the first strand cDNA template synthesized in a reverse transcription reaction on a total RNA template isolated from the roots of *S. altissima*. In these experiments, the first strand cDNA was synthesized using MINT reverse transcriptase (Evrogen, Moscow, Russia) and the (dT)12VN primer.

For the amplification of the 5’- and 3’-end fragments, we used the Encyclo DNA polymerase (No. PK002, Evrogen, Moscow, Russia) and gene-specific primers designed based on the previously identified partial coding sequence of the *NAR2* family gene from *S. altissima* [62]. The 5’-end of *SaNAR2* cDNA was amplified with primers SaNRT3.2_R (round 1) and SaNRT3.2_R1 (round 2). The 3’-end of *SaNAR2* cDNA was amplified with SaNRT3.2_F (round 1) and SaNRT3.2_F1 (round 2). The primers Mix1 (round 1) and Mix2 (round 2) from the manufacturer kit were also used for amplification of *SaNAR2* fragments. Using the Quick-TA kit (No. TAK02, “Evrogen”, Moscow, Russia), the obtained amplicons were cloned into the pAL2-T vector for replication in *Escherichia coli* cells (strain XL1 Blue) and sequenced.

Overlapping sequences of the 5’- and 3’-ends of *SaNAR2* coding sequence were in silico aligned with each other using SnapGene Viewer software, v. 5.0.8 (https://www.snapgene.com/snapgene-viewer). The resulting sequence obtained in this way contained an open reading frame of 591 bp. Based on this sequence, the gene-specific primers SaNRT3.2_F3 and SaNRT3.2_R3 were selected, complementary to the 5’- and 3’- ends of the full-length coding sequence *SaNAR2*. The full-length cDNA of *SaNAR2* was amplified using these primers and the Encyclo DNA polymerase (PK002, Evrogen, Moscow, Russia) for the first 15 cycles, and the ready-to-use CloneAmp HiFi PCR Premix (No. 639298, Clontech, USA) for the next 30 cycles. An amplicon of 591 bp was obtained, named *SaNAR2.2* (see Results section), which was further cloned into the yeast integrative plasmid (shuttle vector) pCCUR2 under the control of the inducible nitrate reductase (NR) promoter *pYNR1* and terminator *tYNR1* of *H. polymorpha*. Promoter *pYNR1* and terminator *tYNR1* sequences were amplified from the *H. polymorpha* genomic DNA template using primer pairs pCCUR2_pYNR1_F and pYNR1_SaNRT3.2_R, tYNR1_SaNRT3.2_F and pCCUR2_tYNR1_R. The first 10 cycles of amplification of the promoter and terminator were performed using Encyclo polymerase (No. PK002, Evrogen, Moscow, Russia); the next 25 cycles were performed using CloneAmp HiFi PCR Premix kit (No. 639298, Clontech, Mountain View, CA, USA). The pCCUR2 plasmid was linearized in the KpnI restriction site and ligated with the synthesized *pYNR1*, *tYNR1* and *SaNAR2.2* sequences using a Gibson assembly kit (No. E5510, SkyGen, NEB, Ipswich, MA, USA) to produce the pCCUR2-*pYNR1-SaNAR2.2-tYNR1* construct (further denoted as pCCUR2- SaNAR2.2). The bacterial *E. coli* XL1 Blue strain was used for routine plasmid propagation. The cloned sequence *SaNAR2.2* (591 bp) was verified by sequencing (“Evrogen”, Moscow, Russia).

### 2.7. Quantitative analysis of SaNAR2.2 transcripts in S. altissima organs

Quantitative analysis of *SaNAR2.2* transcripts in *S. altissima* organs was performed by quantitative real-time PCR (qRT-PCR) using the LightCycler® 96 System (Roche Diagnostics Corporation, Indianapolis, IN, USA). The cDNA samples for the analysis of *SaNAR2.2* expression were synthesized on templates of the total RNA preparations isolated from the organs (roots, leaves, stems) of *S. altissima* plants, grown at various concentrations of nitrate and NaCl in the medium.

PCR analysis was performed using a ready-to-use reaction mixture containing the intercalating dye SYBR Green I (Evrogen, Moscow). As a reference gene, the elongation factor 1 alpha gene from the *S. altissima*, SaeEF1alpha (GenBank ID: MN076325.1), was used. The PCR reaction included 50 ng of cDNA in a total volume of 20 μL. The reactions were initially denatured (95°C/300 s), then subjected to 45 cycles of 95°C/20 s, 58°C/30 s, 72°C/15 s. The results were processed by LightCycler 96SW 1.1 software. The results obtained in these experiments are based on three biological and three analytical replicates. Primer specificity for each gene was evaluated by analyzing the melting curves of PCR products.

*2.8. Bimolecular fluorescence complementation (BiFC) in Nicotiana benthamiana leaves* Plasmid vector constructs for analyzing the interaction of SaNAR2.2 protein with NRT2 family proteins from *S. altissima*, SaNRT2.1 and SaNRT2.5, were obtained based on the pSPYNE-35S and pSPYCE-35S plasmids [63, 64]. The plasmids contain, respectively, the N-terminal fragment (155 aa) and the C-terminal fragment (84 aa) of the fluorescent protein mVenus under the control of constitutive cauliflower mosaic virus 35S promoter. The sequence *SaNAR2.2* was subcloned into the pSPYCE-35S plasmid upstream of the C-terminal fragment of mVenus, and the resulting construct was designated as pSPYCE-35S- SaNAR2.2. The sequences *SaNRT2.1* (OR909030.1) or *SaNRT2.5* (OR828748.1) [48] were cloned into the pSPYNE-35S plasmid upstream of the sequence encoding the N-terminal fragment of mVenus. Accordingly, the constructs pSPYNE-35S-SaNRT2.1 and pSPYNE-35S-SaNRT2.5 were obtained. The obtained constructs were transferred into *Agrobacterium tumefaciens*, strain GV3101. The transformation of bacteria was carried out using the freeze-thaw method [65]. The transformed bacteria were used for the transient transformation of epidermal cells of *N. benthamiana* leaves. Bacterial transformants, as well as the helper strain *A. tumefaciens* containing the coding sequence for the protein p19, were induced with acetosyringone for 3 hours [66]. Suspensions of transformants carrying plasmids with transgenes from *S. altissima* were adjusted to 0.7 OD600, while the suspension of the helper strain was adjusted to 1.0 OD600. Suspensions of the helper strain, the transformant carrying *SaNAR2.2*, and the transformant carrying *SaNRT2.1/SaNRT2.5* were mixed in equal volumes, and the resulting mixture was used to infiltrate the leaves of forty-day-old *N. benthamiana* plants at the abaxial side. After infiltration, the plants were incubated at 21°C for 3 days for transgene expression, after which the leaf fragments were examined with a laser scanning confocal microscope LSM-710-NLO (Carl Zeiss, Jena, Germany). The images of the fluorescence signal from mVenus in *N. benthamiana* cells transformed with the vector constructs were obtained at λex 515 nm and λem 527 nm. The transmitted laser light was recorded with a separate T-PMT detector.

### 2.9. Complementation assay with pCCUR2-SaNAR2.2 and pCHLX-SaNRT2.1/pCHLX- SaNRT2.5 constructs

To study the function of SaNAR2.2 protein and the SaNRT2.1/SaNRT2.5 proteins, the mutant yeast strain *Δynt1* was co-transformed with pairs of integrative vector constructs: pCHLX-SaNRT2.1/pCCUR2, or pCHLX-SaNRT2.5/pCCUR2, or pCHLX- SaNRT2.1/pCCUR2-SaNAR2.2, or pCHLX-SaNRT2.5/pCCUR2-SaNAR2.2. The constructs pCHLX-SaNRT2.1 and pCHLX-SaNRT2.5 were obtained by us earlier [48]. In these constructs coding sequences *SaNRT2.1* and *SaNRT2.5* were under the control of the inducible nitrate reductase (NR) promoter *pYNR1* and terminator *tYNR1* of *H. polymorpha*. Wild type strain DL-1 and the mutant strain *Δynt1* co-transformed with vector pair pCHLX/pCCUR2 were taken as controls. Vector constructs were linearized at the PspEI site (for pCHLX and pCHLX-SaNRT2.5), EgeI site (for pCHLX-SaNRT2.1), or PspCI site (for pCCUP2 and pCCUR2-SaNAR2.2) and *H. polymorpha* cells were transformed with the obtained linear plasmid forms. Yeast cells were transformed by electroporation [67] using an Eppendorf device (Eppendorf, Framingham, MA, USA). Yeast transformants were selected on minimal selective synthetic SD media in the absence of leucine and uracil. Selected yeast transformants were grown in liquid SD medium containing 0.5% ammonium sulfate overnight at 37◦C. Then, 1 ml of the yeast cultures were centrifuged for 5 min at 2500g, the cell precipitates were washed with sterile water, resuspended in 1 ml water and 2 μL each the obtained suspensions were placed on agarized SD medium containing KNO_3_ (0.2, 0.5, 1.0, 2.5, and 5.0 mM) instead of ammonium sulfate. The plates were incubated at 37◦C for 2–3 days until colonies appeared.

### 2.10. Vector constructs pCCUR2-SaNAR2.2-mCherry and pCHLX-SaNRT2.5-GFP

For the expression of SaNAR2.2 fused at the C-terminus with red fluorescent protein mCherry and SaNRT2.5 fused at the C-terminus with green fluorescent protein GFP in *H. polymorpha* cells, the pCCUR2-SaNAR2.2-mCherry and pCHLX-SaNRT2.5-GFP constructs were obtained where coding sequences *SaNAR2.2::mCherry* and *SaNRT2.5::GFP* were under the control of the inducible nitrate reductase promoter *pYNR1* and terminator *tYNR1* of *H. polymorpha*. GFP coding sequence was amplified from plasmid pTR-UF12 [68] with GFP_Gib_F and tYNR1_GFP_R primers. mCherry coding sequence was amplified from pcDNA3.1/hChR2(H134R)-mCherry plasmid with mCherry_Gib_F and tYNR1_mCherry_R primers. pCHLX-SaNRT2.5 and pCCUR2-SaNAR2.2 constructs were linearized by inverse PCR with corresponding primers. Ligation of fluorescent protein coding sequences with linearized constructs was carried out using Gibson assembly kit (NEB, Ipswich, MA, USA). Final coding sequences of interest in pCHLX and pCCUR2 vectors were verified by sequencing.

### 2.11. Bioinformatics analysis

The virtual translation of the nucleotide sequence *SaNAR2.2* into an amino acid sequence was carried out using the online service on the ExPASy portal (http://web.expasy.org/translate/). The molecular weight of SaNAR2.2 protein was also determined using the service on this portal (http://web.expasy.org/protparam/). The cellular localization of the protein was predicted using the on-line service WoLF PSORT II on the GenScript portal (https://www.genscript.com/wolf-psort.html). The affiliation of SaNAR2.2 protein with the NAR2/NRT3 family and the prediction of its functions were carried out using the InterPro v.102.0 resource (http://www.ebi.ac.uk/interpro/) as well as the Protein BLAST algorithm on the NCBI portal (http://www.ncbi.nlm.nih.gov/). The transmembrane topology of the protein was predicted using on-line DeepTMHMM software v.1.0.24 (https://dtu.biolib.com/DeepTMHMM). The 3D-structure models and the model of interaction between SaNAR2.2 and SaNRT2.5 proteins was created using AlfaFold3 software [69] at alphafoldserver.com/.

Multiple alignment of amino acid sequences of NAR2 family proteins was performed online using the MAFFT v.7 software (https://www.ebi.ac.uk/Tools/msa/mafft/) and visualized using Jalview v. 2.11.2.7 software (https://www.jalview.org/). Phylogenetic analysis of NAR2 family proteins was performed using the Molecular Evolutionary Genetic Analysis software (MEGA v.11; https://www.megasoftware.net/), employing the maximum likelihood method based on the Jones–Taylor–Thornton model [70] (1000 bootstrap replicates were performed). Amino acid sequences of NAR2 family proteins for comparative analysis were extracted from the NCBI portal (https://www.ncbi.nlm.nih.gov/protein/).

## Results

### 3.1. Cloning of the full-length coding sequence of SaNAR2.2 and bioinformatic analysis of the SaNAR2.2 protein

At earlier stages of our search for sequences encoding nitrate-transporting proteins of the halophyte *S. altissima*, a partial coding sequence (CDS) of the *NAR2* family gene from *S. altissima* was identified [62]. In the present study, gene-specific primers based on the previously identified partial sequence were designed, and the 3’- and 5’-end fragments of the full-length CDS of the *NAR2* family gene from *S. altissima* were amplified using the Step-Out RACE method (Fig. 1a, b); then the full-length targeted cDNA of 591 bp was amplified (Fig. 1c). For amplification of the target sequence, the first strand cDNA was synthesized from total RNA extracted from the roots of *S. altissima* was used, as it is known since *NAR2* genes are predominantly expressed in plant roots [20, 39]. The 591 bp full-length CDS of *SaNAR2* was cloned into yeast integrative plasmid pCCUR2 for further expression in the cells of the methanol-utilizing yeast *H. polymorpha* (see below). The cloned sequence has been deposited in GenBank (ID: OR828749.1).

**Figure 1.**
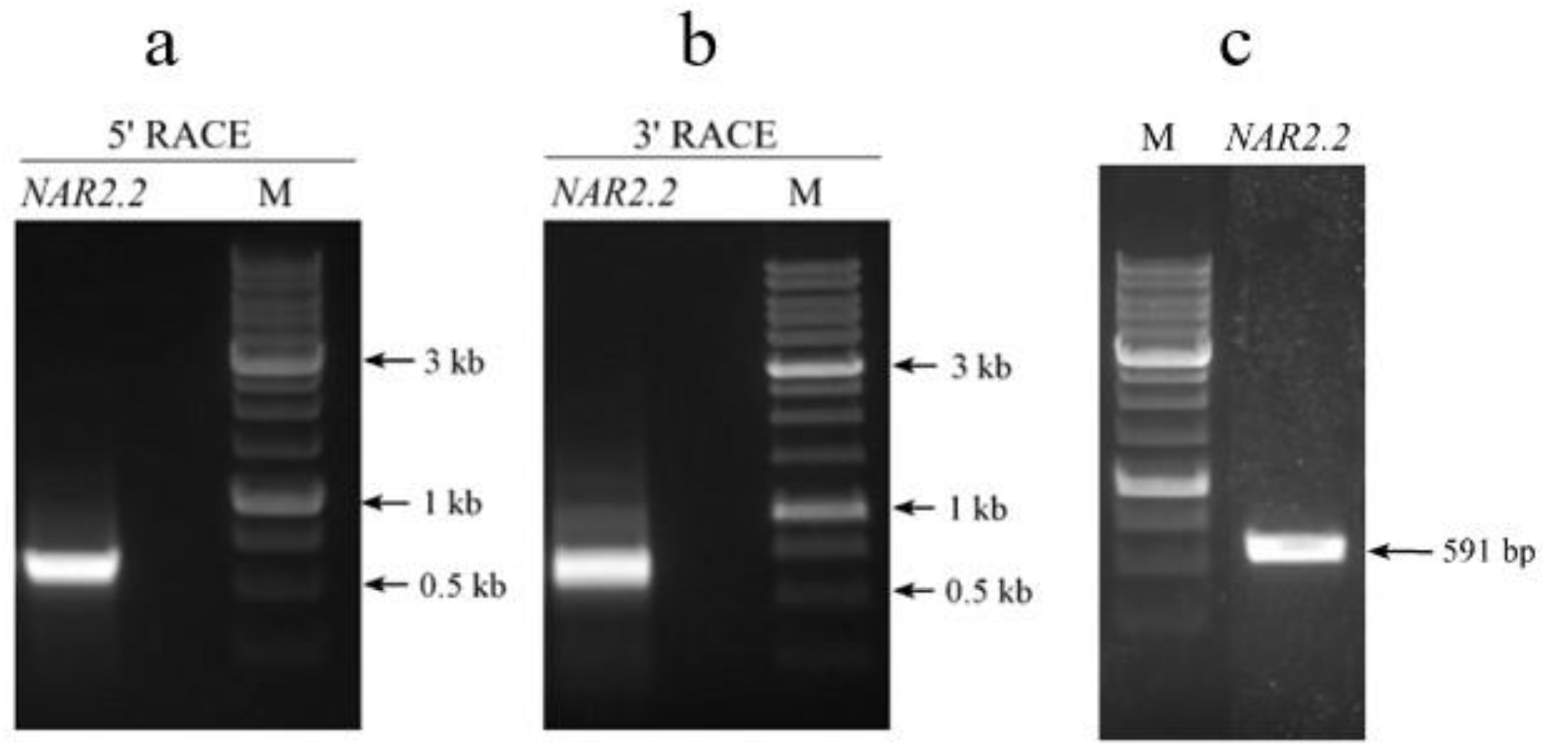
(**a**, **b**): Analysis of 5′-end and 3′-end fragments of *SaNAR2.2* coding sequence synthesized on the total cDNA template from *S. altissima* roots using Step-Out RACE technology. (**c**): Analysis of the full-length *SaNAR2.2* coding sequence that was used for further cloning in plasmid pCCUR2. DNA fragments were separated by electrophoresis in 1% agarose gel. M —DNA molecular weight markers.

Sequencing results showed that the cloned *NAR2* sequence from *S. altissima* contained an open reading frame of 591 bp for a protein of 196 amino acids in size, with a calculated molecular mass of 21.7 kDa (GenBank ID: WPH61291.1). The affiliation of this protein with the NAR2/NRT3 protein family, i.e., proteins involved in high-affinity nitrate transport in plants, was confirmed using the online resource InterPro. On the phylogenetic tree of NAR2 family proteins, the protein from *S. altissima* lays in a clade with the proteins of the NAR2.2 subfamily from other plants of Amaranthaceae family (Fig. 2a); therefore, we classified it into this subfamily and named it SaNAR2.2.

**Figure 2.**
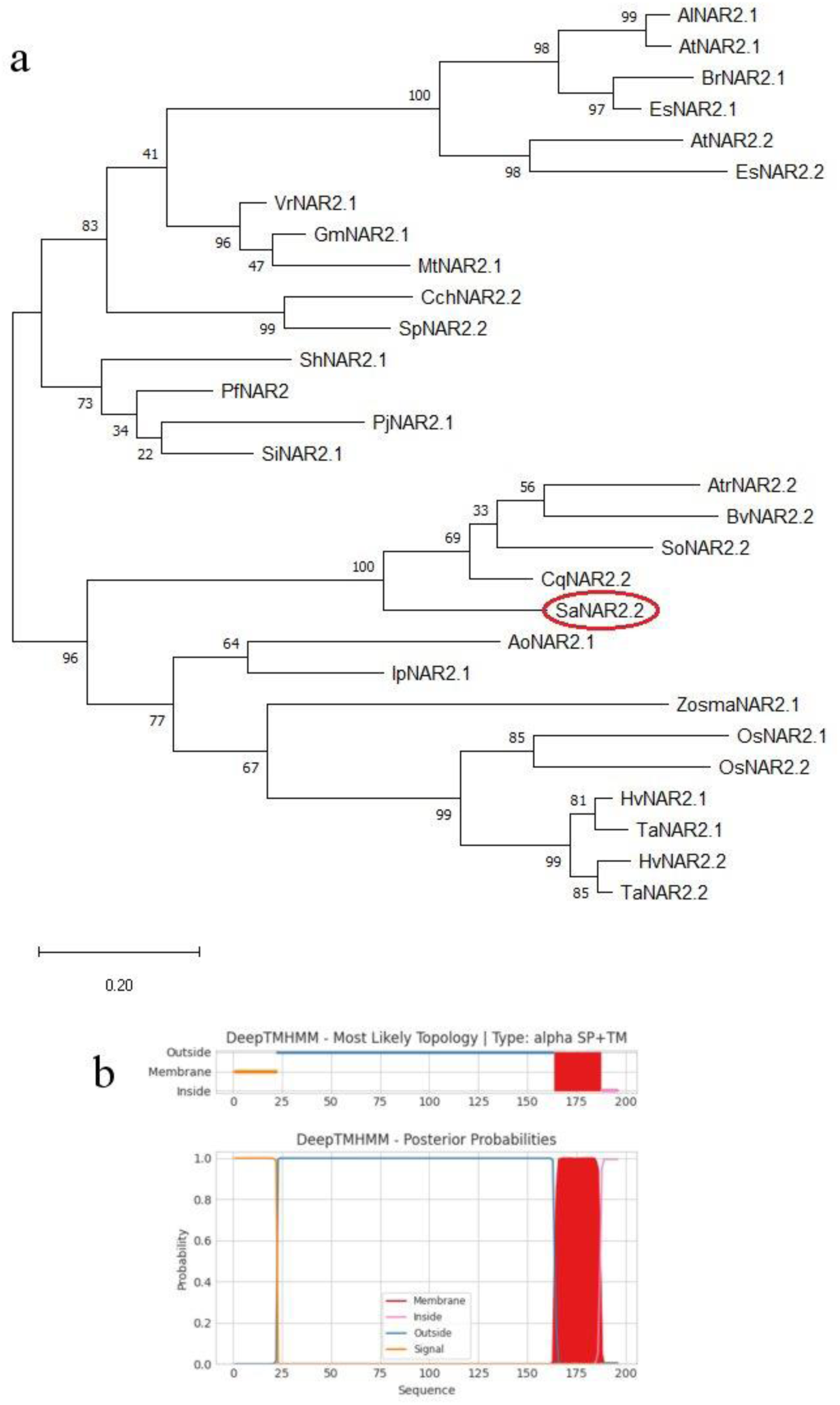
(a) An unrooted phylogenetic tree of plant NAR2 proteins including the SaNAR2.2 protein of *S. altissima*. The phylogenetic tree was built in MEGA v.11 using the maximum likelihood method based on the Jones–Taylor–Thornton model. The number of bootstrap replicates was 1000; the values of bootstrap support are indicated near the nodes. The NAR2 protein sequences were extracted from the NCBI portal (https://www.ncbi.nlm.nih.gov/protein/, accessed on 15 October 2024). Names of plant species and protein IDs are given in Table S2 (Supporting information). (**b**) Two-dimensional topology of SaNAR2.2 protein predicted using DeepTMHMM software (v.1.0.24). SaNAR2.2 has one transmembrane segment in the C-terminal region (164-187 aa). The large N-terminal part of the SaNAR2.2 is located in the extracellular space. A signal sequence (1-22 a.a.) is also present at the N-end.

According to the prediction of the cellular localization and protein topology, SaNAR2.2 is a resident of the plasma membrane; it has one transmembrane segment at the C-terminal region (164 – 187 aa), and a small C-end (188 – 196 aa) located in the cytoplasm. The large N-terminal part of the SaNAR2.2 protein (1 – 163 aa) is located in the extracellular space (Fig. 2b). A signal sequence (1-22 aa) was also identified at the N-terminus.

The alignment of the amino acid sequence of SaNAR2.2 protein with the sequences of some other NAR2 proteins showed that proteins of this family from various plant sources possess conserved domains in the central protein part and in the C-terminal region (Fig. 3). This data suggested the involvement of the amino acid sequences of these regions in interactions with partner proteins of the NRT2 family.

**Figure 3.**
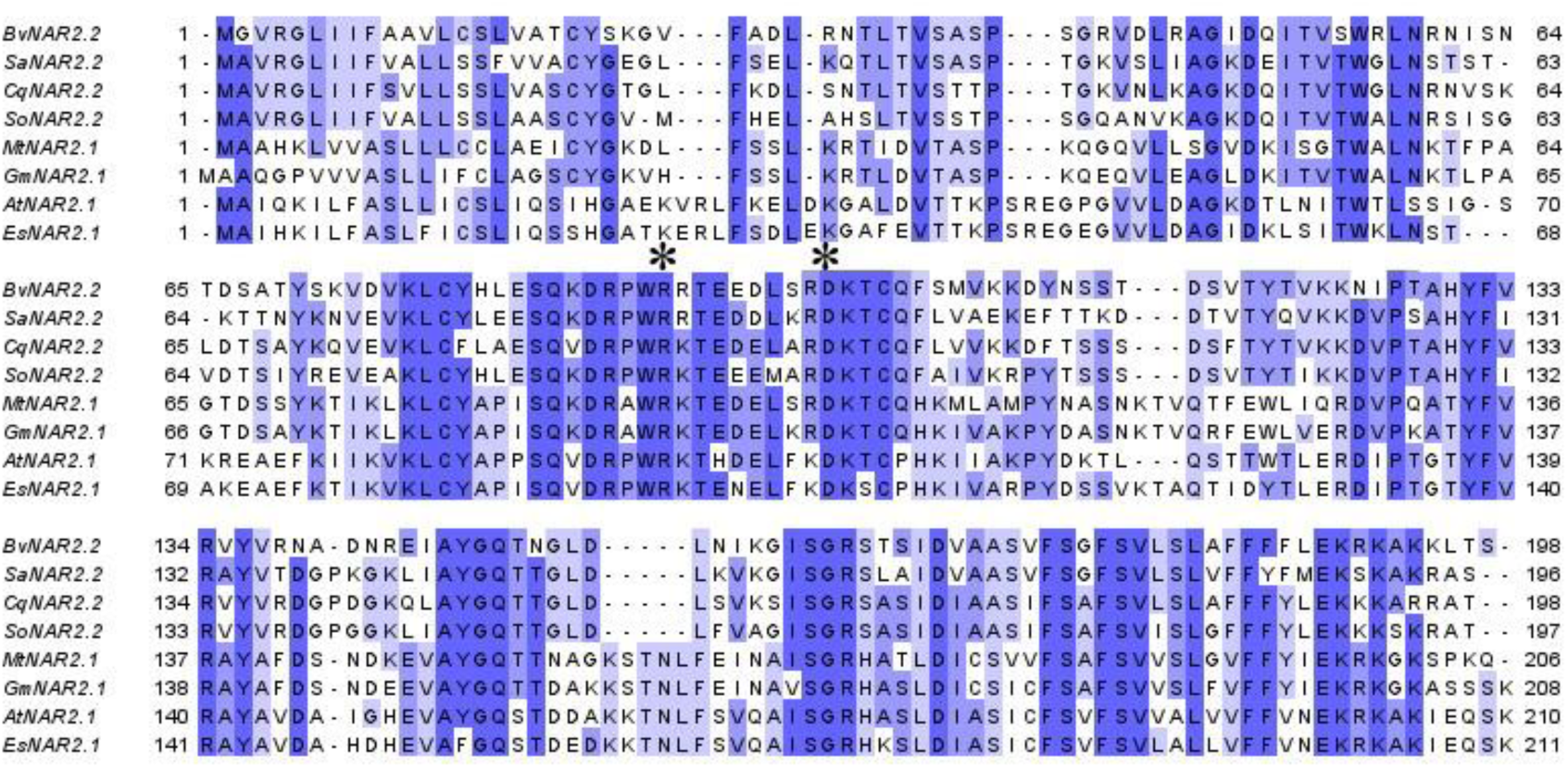
Multiple sequence alignment performed in Clustal Omega software (https://www.ebi.ac.uk/jdispatcher/msa/clustalo, accessed on 22 October 2024) for NAR2 proteins from *Arabidopsis thaliana* (AtNAR2.1: NP_199831.1), *Beta vulgaris* (BvNAR2.2: XP_010667500.1), *Chenopodium quinoa* (CqNAR2.2: XP_021720161.1), *Eutrema salsugineum* (EsNAR2.1: XP_006402168.1), *Glycine max* (GmNAR2.1: XP_003549822.1), *Medicago truncatula* (MtNAR2.1: XP_013457733.1), *Spinacia oleracea* (SoNAR2.2: XP_021863523.1), and *Suaeda altissima* (SaNAR2.2: WPH61291.1). Protein GenBank IDs are indicated in the parenthesis. The key amino acid residues which are presumably crucial for interaction with the transporters of NRT2 family are marked by asterisks. The intensity of the staining of amino acid residues corresponds to the degree of their identity (percentage identity).

Indeed, it has been shown that in central region, NAR2 proteins have important conserved residues of arginine R and aspartate D, which are crucial for interaction with NRT2 family transporters [40, 71]. The location of these residues in different NAR2 proteins varied in different species. For example, these are R96 and D105 for the AtNAR2.1 from *A. thaliana* [71] and R100 and D109 for the OsNAR2.1 from rice [40]. In the central region of the SaNAR2.2, these conserved arginine and aspartate residues apparently correspond to the amino acid residues R88 in the conserved motif WR(K, R) and D97 in the motif RDKTCQ (Fig. 3). The modeling of the three-dimensional structures of SaNAR2.2 (Fig. 4a), SaNRT2.5 (the high affinity nitrate transporter from *S. altissima*, presumably a partner protein for SaNAR2.2) (Fig. 4b), and the heterodimer of these proteins (Fig. 4c) also predicted that the central region of SaNAR2.2 is involved in interaction with the protein SaNRT2.5. Moreover, the predicted 3D model of the heterodimer demonstrated that the transmembrane C-terminal region of the SaNAR2.2 protein and small C-end domain located in cytoplasm, which form an α-helix (164-196 aa), may also be involved in this interaction (Fig. 4c).

**Figure 4.**
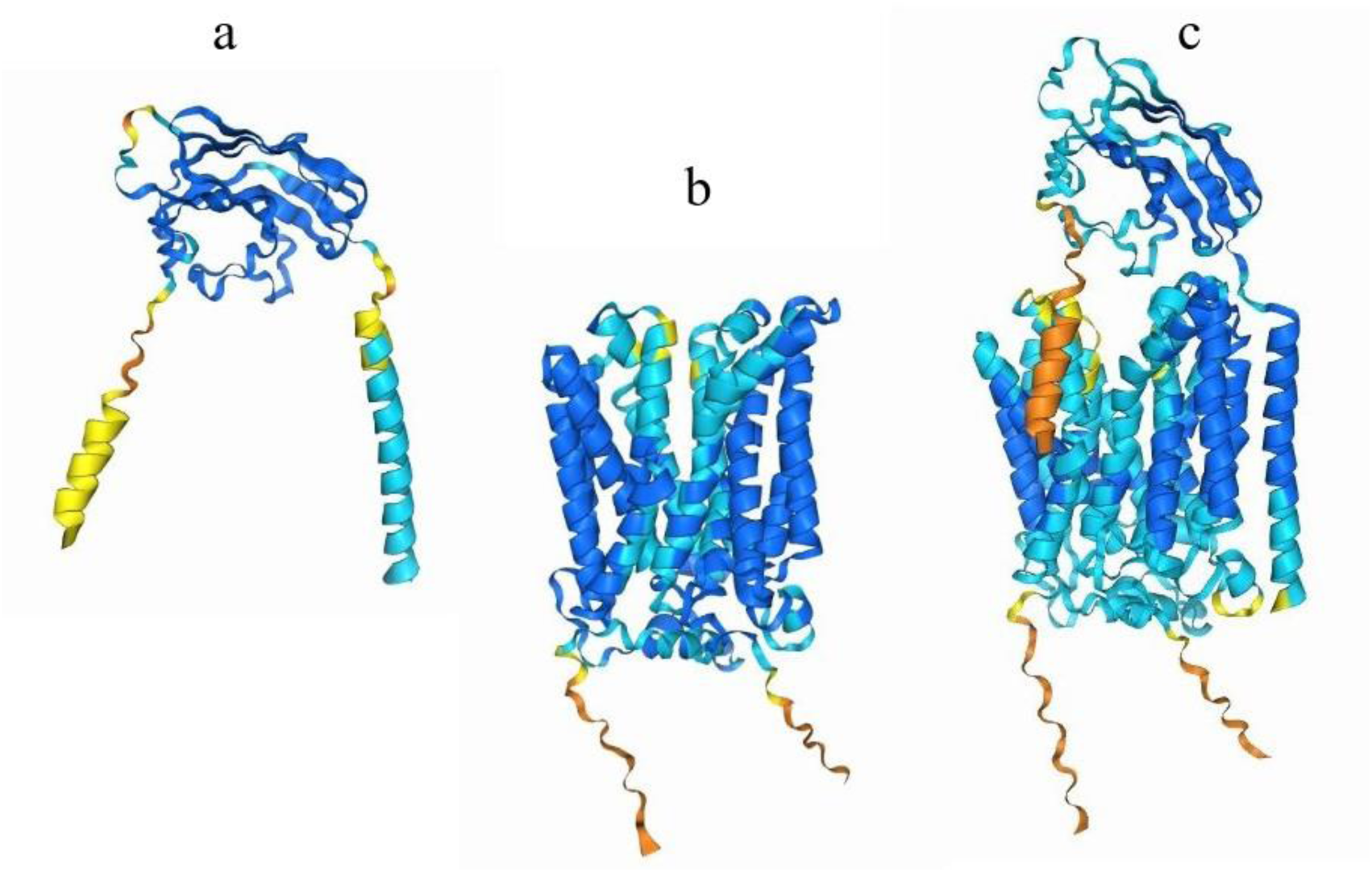
Three-dimensional structure modeling of (**a**) SaNAR2.2 protein, (**b**) SaNRT2.5 protein, and (**c**) SaNAR2.2/SaNRT2.5 heterodimer predicted by AlfaFold3 software (alphafoldserver.com/). The model predicts that the central region of SaNAR2.2 and the transmembrane C-terminal region which forms an α-helix are involved in interaction with the protein SaNRT2.5.

### 3.2. Detection of SaNAR2.2 and SaNRT2.1/SaNRT2.5 protein interaction

The question of whether the proteins SaNAR2.2 and SaNRT2.1/SaNRT2.5 interact was clarified using the BiFC method in *N. benthamiana* leaves. Co-expression of SaNAR2.2 fused with the C-terminal fragment of the yellow fluorescent protein mVenus and SaNRT2.1 or SaNRT2.5, each fused with the N-terminal fragment of mVenus, resulted in the recovery of mVenus fluorescence (Fig. 5a, b). The latter demonstrated that two nonfluorescent N- and C-terminal halves of the fluorescent protein were brought together through the interaction of SaNAR2.2 and SaNRT2.1 or SaNRT2.5.

**Figure 5.**
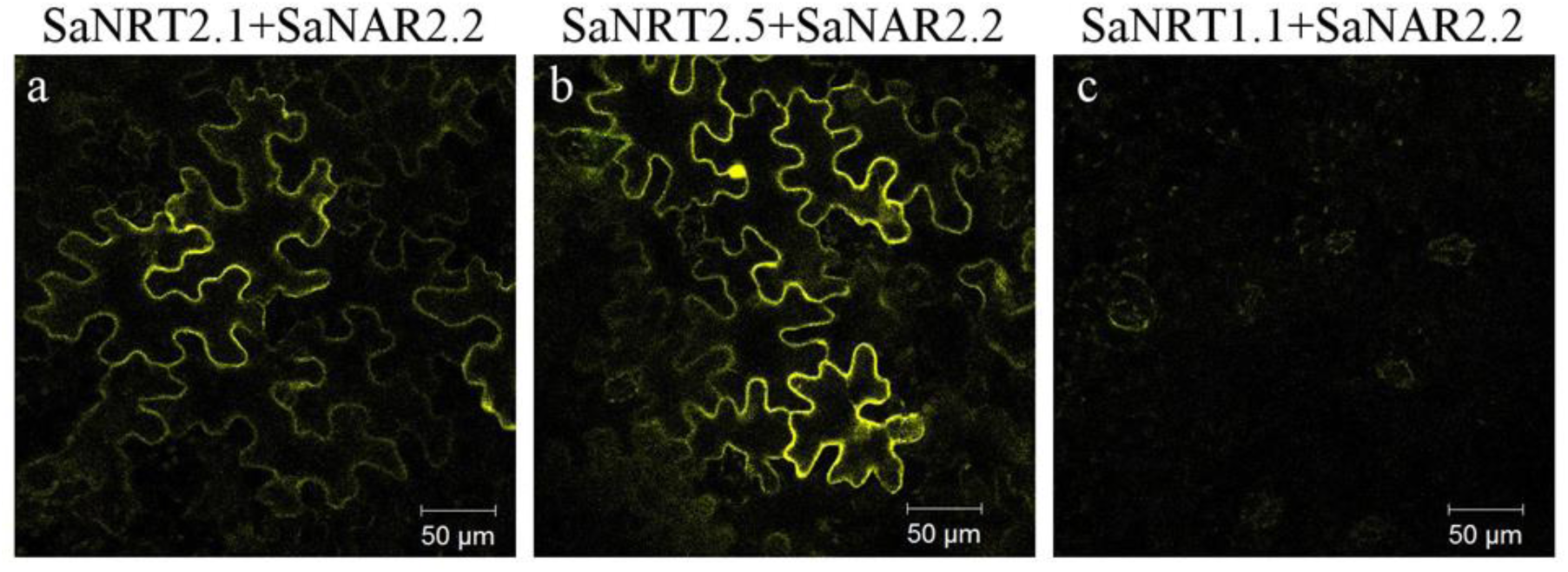
Detection of the SaNRT2.1/SaNAR2.2 (a) or SaNRT2.5/SaNAR2.2 (b) interactions in leaves of *N. benthamiana* plants using BiFC method. Transient co-expression of *SaNAR2.2* fused with the C-terminal half of the fluorescent protein mVenus and *SaNRT2.1* or *SaNRT2.5*, each fused with the N-terminal half of mVenus, in *N. benthamiana* leaves resulted in the recovery of mVenus fluorescence (a, b), which could be observed at the periphery of transformed tobacco cells. The images of the fluorescence signal were obtained with a laser scanning confocal microscope LSM-710-NLO (Carl Zeiss, Jena, Germany) at λex 515nm and λem 527nm. In these experiments, *SaNRT1.1*, a low-affinity nitrate transporter gene of the *NRT1* family from *S. altissima* [57], was co-expressed with SaNAR2.2 in the leaves of *N. benthamiana* as a negative control (c). Each confocal image is a representative picture of eight independent assays.

The fluorescence of mVenus could be observed at the periphery of transformed tobacco cells, which indicated localization of the interacting proteins SaNRT2/SaNAR2.2 presumably at the plasma membrane. In these experiments, *SaNRT1.1* (GenBank ID: OQ330855.1), a low-affinity nitrate transporter gene of the *NRT1* family from *S. altissima* [57], was expressed in the leaves of *N. benthamiana* as a negative control with the pSPYNE-35S-SaNRT1.1 construct, carrying *SaNRT1.1* sequence fused with the N-terminal fragment of mVenus. Since NAR2 family proteins interact with NRT2 proteins but not with NRT1 proteins, no mVenus protein fluorescence was observed in tobacco leaves co-transformed with pSPYCE-35S-SaNAR2.2 and pSPYNE-35S-SaNRT1.1 constructs (Fig. 5c).

### 3.3. Expression of SaNAR2.2 in S. altissima organs under various ionic conditions

The expression of *SaNAR2.2* gene in the organs of *S. altissima* plants, grown under low (0.5 mM) and high (15 mM) nitrate concentrations in the nutrient medium in the absence of NaCl or in the presence of increasing salt concentrations (250 mM, 500 mM, 750 mM NaCl), was analyzed using qRT-PCR method. In these experiments, the elongation factor 1 alpha gene from *Suaeda* was used as the reference gene (SaeEF1alpha, GenBank ID: MN076325.1). Earlier, we showed that the expression of this housekeeping gene was constitutive in various *Suaeda* organs and under different experimental conditions [72].

It was found that *SaNAR2.2* gene was expressed nearly exclusively in roots; in leaves and stems the gene expression level was negligible (Fig. 6a). In the absence of salt in the medium, the expression of *SaNAR2.2* was at comparable levels in the roots of plants grown at low (0.5 mM) or high (15 mM) NO_3_^̶^ concentrations; with the increase in salt concentration in the medium, the expression of *SaNAR2.2* decreased provided the nutrient medium contained high concentrations of nitrate, and conversely, the gene expression significantly increased at low nitrate concentrations in the nutrient medium, reaching maximum values at 500 mM NaCl. At this salt concentration, the gene expression level increased almost tenfold (Fig. 6b). Further increase of NaCl concentration in the nutrient solution to 750 mM led to decrease in the expression of *SaNAR2.2* gene. Likewise, under these conditions, i.e. low nitrate content and increasing NaCl concentrations, the expression of *SaNRT2.1* and *SaNRT2.5* (which encode the potential partners of the SaNAR2.2 protein, high-affinity nitrate transporters of the NRT2 family in *S. altissima,* and whose expression also occurs predominantly in *S. altissima* roots [48]), changed in a similar way (Fig. 6 c, d). However, the expression levels of these three genes, *SaNAR2.2*, *SaNRT2.1*, and *SaNRT2.5*, differed significantly. The expression of *SaNAR2.2* reached an extremely high level, almost twice that of the house-keeping gene *SaeEF1ɑlfa*, whereas the expression levels of *SaNRT2.1* and *SaNRT2.5* were significantly lower.

**Figure 6.**
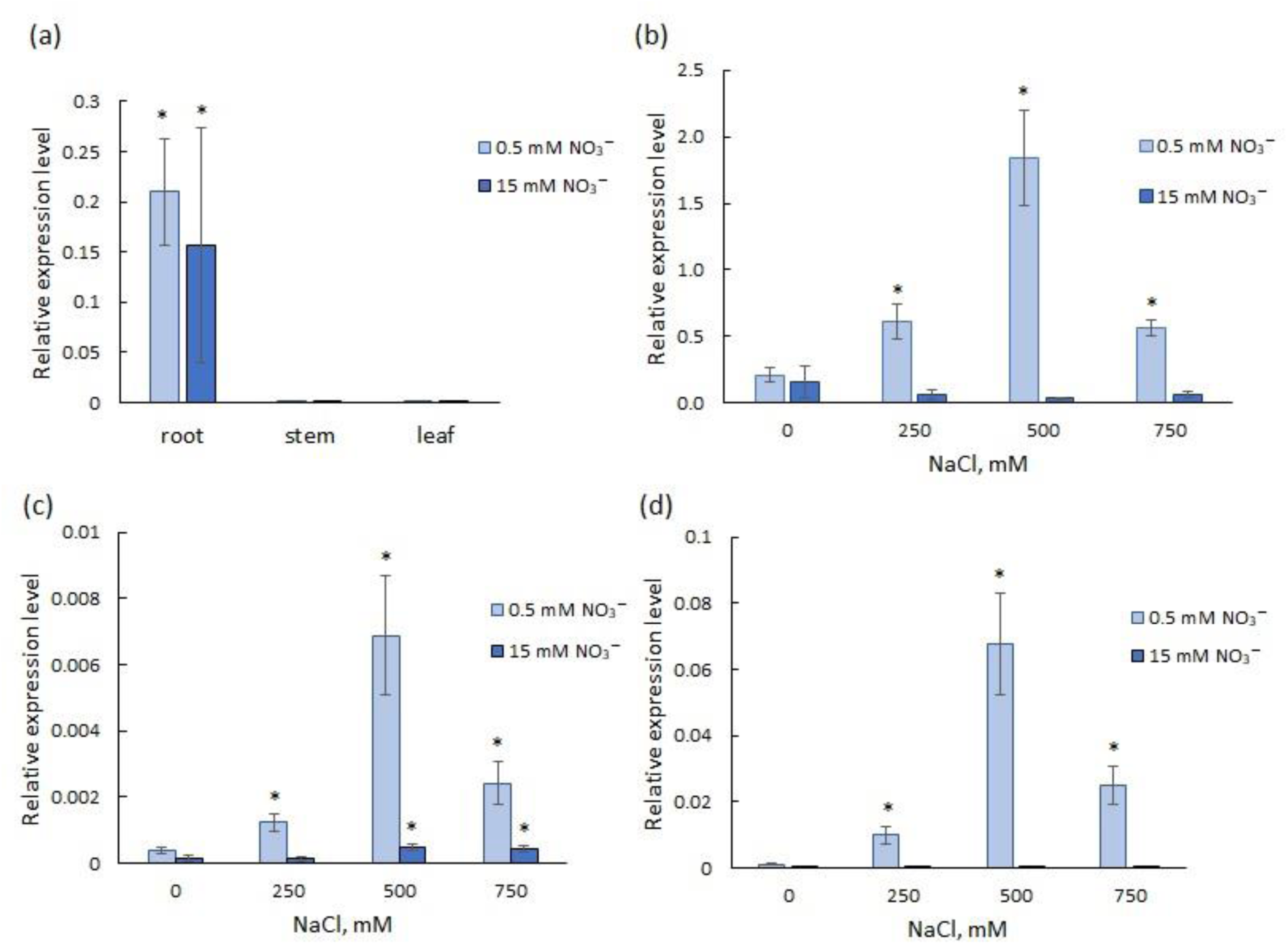
Relative abundance of *SaNAR2.2* transcripts (a, b), *SaNRT2.1* transcripts (c), and *SaNRT2.5* transcripts (d) in the roots of *S. altissima* plants grown in the nutrient medium containing low (0.5 mM) or high (15 mM) NO_3_^−^ concentrations under increasing NaCl concentrations. Changes in expression relative to the reference gene are presented, the Sae*EF1ɑ* gene from *S. altissima* was used as a reference gene. A *p*-value < 0.05 was considered to be statistically significant. * *p* < 0.05. Standard deviations are given.

### 3.4. Complementation analysis of SaNAR2.2 function

To determine the possible functional role of SaNAR2.2 in complex with SaNRT2.1 or SaNRT2.5 proteins in NO_3_^̶^ transport, the cells of the yeast *H. polymorpha* mutant strain *Δynt1*, which lacks the only gene for the nitrate transporter in this organism, were transformed with plasmids carrying *SaNRT2.1* or *SaNRT2.5* coding sequences, or co-transformed with the pair of plasmid constructs carrying *SaNRT2.1*/*SaNRT2.5* and *SaNAR2.2* coding sequences. All the yeast strains were plated on agarized nutrient media containing a range of nitrate concentrations (Figure 7). The growth of mutant strain *Δynt1* on the media containing nitrate as a nitrogen source was expectedly suppressed. Unfortunately, we did not see a clear picture of the recovery of growth of the mutant *Δynt1* transformed with vector constructs carrying *SaNRT2.1* or *SaNRT2.5* sequences or co-transformed with two plasmids carrying SaNRT2.1/SaNRT2.5 and SaNAR2.2 on these media. The growth of yeast transformants was significantly weaker than the growth of wild-type cells with an intact nitrate uptake mechanism.

**Figure 7.**
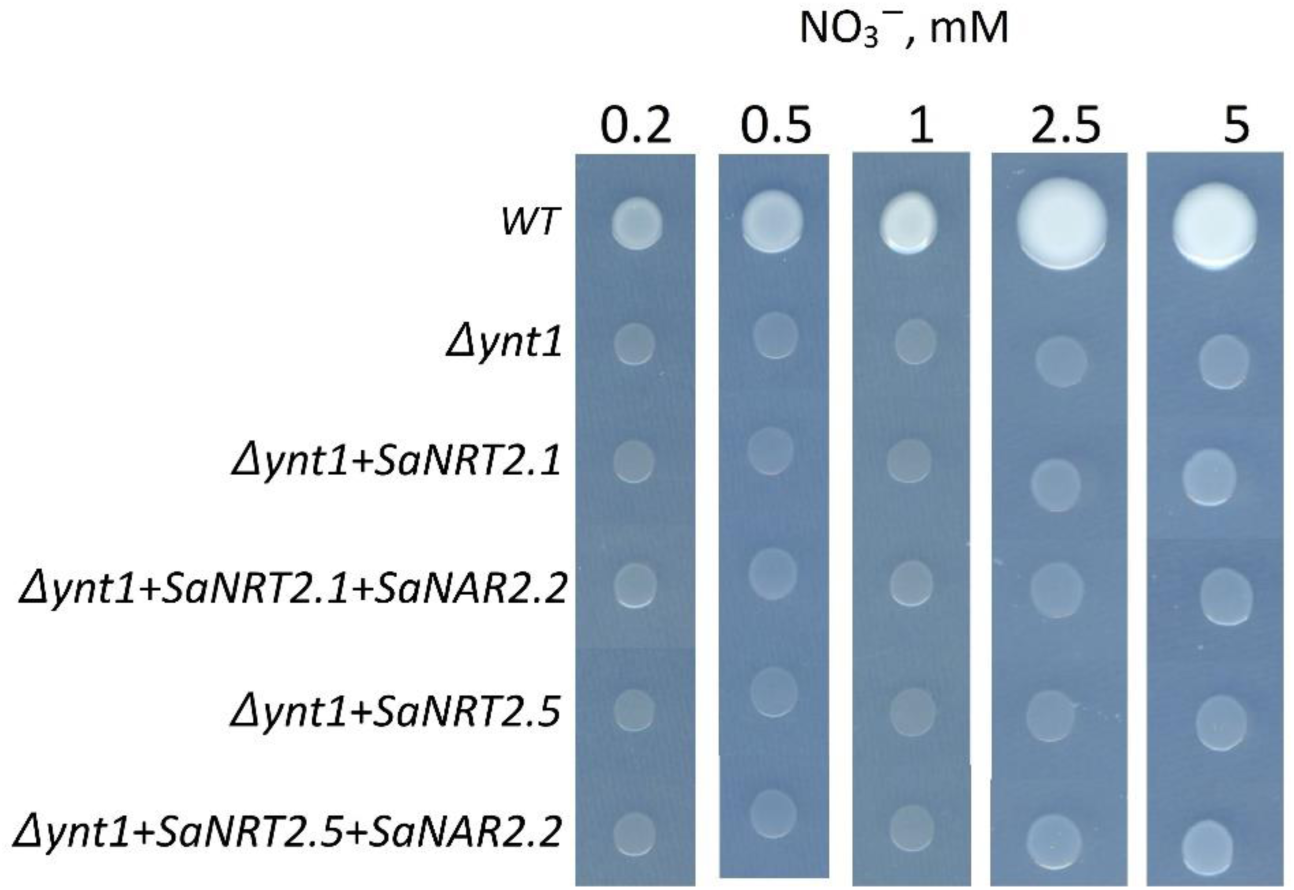
Complementation assay of the *H. polymorpha* mutant strain *Δynt1* co-transformed with pairs of integrative plasmid constructs: pCHLX-SaNRT2.1/pCCUR2, or pCHLX-SaNRT2.5/pCCUR2, or pCHLX-SaNRT2.1/pCCUR2-SaNAR2.2, or pCHLX-SaNRT2.5/pCCUR2-SaNAR2.2. Wild type strain DL-1 and the mutant strain *Δynt1* co-transformed with plasmid pair pCHLX/pCCUR2 were taken as controls. Yeast grew on minimal agarized SD medium supplied with indicated concentrations of NO_3_^−^. Approximately 10^5^ cells of each strain were plated on Petri dishes and incubated at 37 °C for 3 days.

The possible reason for weak functional complementation of the yeast mutant strain *Δynt1* by *S. altissima* proteins could be the insufficient level of the heterologous protein synthesis in yeast cells. To determine the presence of recombinant *Suaeda* proteins in the transformed yeast cells, the mutant *Δynt1* was co-transformed with vector constructs pCCUR2-SaNAR2.2-mCherry and pCHLX-SaNRT2.5-GFP. The plasmids carried coding sequences, accordingly, for SaNAR2.2 fused at the C-terminus with red fluorescent protein mCherry, and for SaNRT2.5 fused at the C-terminus with green fluorescent protein GFP. Inspection of the transformed strain cells with a laser scanning microscope showed that co-expression of *Suaeda* transgenes occurred in yeast cells. However, the proteins of interest were synthesized only in a small portion of the transformed cells; moreover, these proteins remained localized in the internal space of the yeast cell and did not reach their destination, which is the plasma membrane (Figure 8). Potential causes for such a result are discussed in the Discussion section.

**Figure 8.**
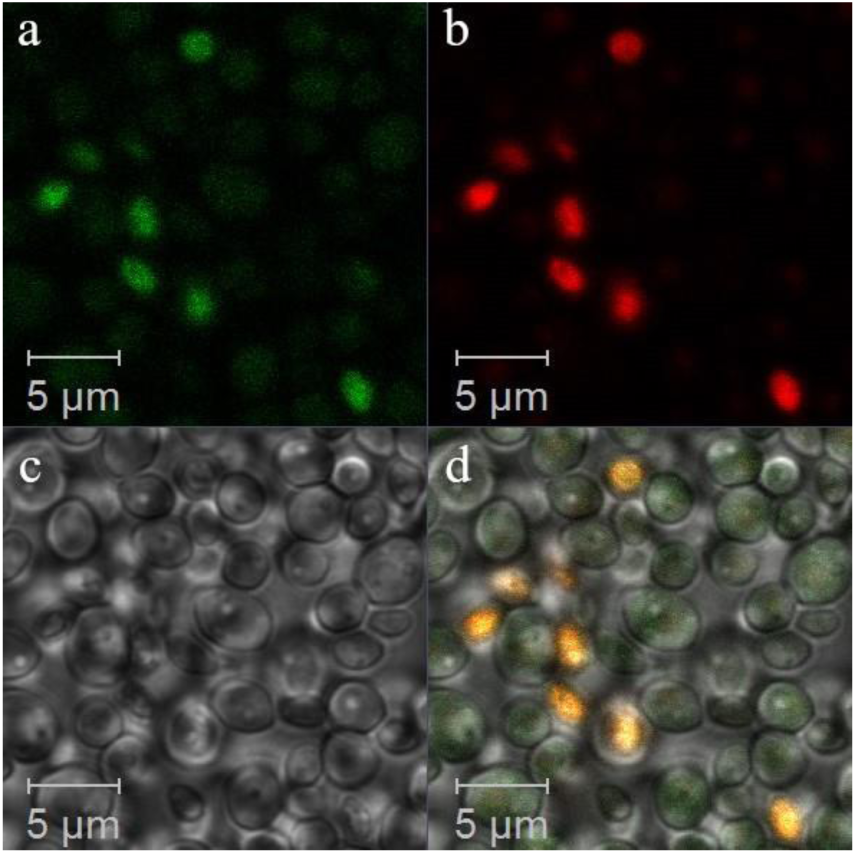
Images of *Δynt1* cells co-transformed with pCHLX-SaNRT2.5-GFP and pCCUR2-SaNAR2.2-mCherry constructs. (a) Green fluorescence of GFP. (b) Red fluorescence of mCherry. (c) Bright field image. (d) Merge of GFP (a) and mCherry (b) fluorescence images (yellow color) and bright field image (c). The images were obtained with a laser scanning confocal microscope LSM-710-NLO (Carl Zeiss, Jena, Germany). The fluorescence images were obtained using excitation/emission wavelength of 488/512 nm for GFP and 587/610 nm for mCherry, respectively.

## Discussion

NAR2 is a family of plasma membrane-associated small proteins with a molecular mass slightly over 20 kDa and a C-terminal transmembrane region. These proteins are an important component of high-affinity nitrate transport system in plants; they act as partners of high-affinity nitrate transporters of NRT2 family [1, 24–26]. Currently available data on high-affinity nitrate uptake systems in plants have been obtained for glycophytes [15, 17, 18, 24, 28–31]. As for salt-tolerant plants, we found the only example of such studies. The high-affinity nitrate transport system ZosmaNRT2.5/ZosmaNAR2 was described in marine seagrass *Zostera marina* [33, 37] while none in land halophytes, in our knowledge.

In our work, the coding sequence of *NAR2* family gene from the euhalophyte *S. altissima* was cloned. The search for the gene of interest in this plant was complicated by the fact that there is no sequenced genome of this species. Therefore, we used an *in silico* search in the de novo assembled transcriptomes of two related halophyte species, *S. fruticosa* and *S. glauca*, where the transcripts of NAR2 family genes were identified by us previously [62]. Transcripts of the only NAR2 gene were found both in *S. fruticosa* and *S. glauca* transcriptomes. The full-length coding sequence *SaNAR2.2* cloned in this work was obtained based on the homology of *NAR2* coding sequences among different *Suaeda* species. Using primers derived from *NAR2* transcripts in *S. fruticosa* and *S. glauca* transcriptomes, we found a single *NAR2* sequence for *S. altissima*. Single *NAR2* gene has also been found in a number of other plants: spinach [18], chrysanthemum [30], maize [32], seagrass *Zostera marina* [33], wheat [34]. It should be noted that even though several *NAR2* family genes have been identified in a plant genome, apparently the only one of NAR2 proteins is involved in nitrate uptake via interactions with NRT2 proteins, as shown for AtNAR2.1 [23, 26] or OsNAR2.1 [29]. Phylogenetic analysis demonstrated that the NAR2 protein from *S. altissima* belongs to the same clade as NAR2.2 proteins from other members of Amaranthaceae family (Fig. 2a), and therefore it was classified to the NAR2.2 subfamily. According to the secondary structure prediction, SaNAR2.2 protein has one transmembrane segment at C-terminal region and a small C-terminal end located in cytoplasm. The large N-terminal part of SaNAR2.2 protein is located in the extracellular space (Fig. 2b).

To detect the interaction of SaNAR2.2 with putative partner proteins from high-affinity nitrate transporter family NRT2 in *S. altissima*, SaNRT2.1 and SaNRT2.5 (previously identified high-affinity nitrate transporters in *S. altissima* [48]), we used BiFC (the bimolecular fluorescence complementation method) and also did *in silico* modelling. The BiFC assay is based on the recovery of fluorescence when two non-fluorescent N-and C-terminal halves of a fluorescent protein are brought together through the interaction of the proteins to which they are fused to [53, 63]. The genetic constructs for analyzing the interaction of SaNAR2.2 protein with SaNRT2.1 and SaNRT2.5 were obtained based on plasmid vectors that encode, respectively, N-terminal half and C-terminal half of the yellow fluorescent protein mVenus. Using the vector constructs, transient expression of the specified *S. altissima* genes was achieved in *N. benthamiana* leaves. Co-expression of SaNAR2.2, fused with C-terminal fragment of mVenus, and SaNRT2.1 or SaNRT2.5, each fused with N-terminal fragment of mVenus, resulted in the recovery of mVenus fluorescence at the periphery of transformed tobacco cells (Figure 5). Obtained results demonstrated that proteins SaNAR2.2 and SaNRT2.1/SaNRT2.5 do indeed interact; the complex of these interacting proteins is most likely located in the plasma membrane, similar to observations with orthologous proteins from other plants [38–40].

The question about essential amino acid residues in NAR2 proteins which are involved in interaction with NRT2 proteins is of great scientific interest. However, so far a very limited number of research studies have investigated this research area issue. For AtNAR2.1/AtNRT3.1 proteins it was demonstrated that the replacement of aspartate D105 in the central part of the protein markedly reduced nitrate uptake and accumulation in *Arabidopsis* plants [77]. Also, for protein OsNAR2.1, it was shown that mutations R100G and D109N disrupted the interaction of this protein with OsNRT2.3a [40]; it indicated that these amino acid residues are important not only for high-affinity nitrate transport activity but also for co-localization of OsNAR2.1 with OsNRT2.3a at the plasma membrane. These arginines and aspartates, which are at a similar distance of nine aa residues from each other, appear to be conservative ones and can be found in the central parts of all NAR2 proteins aligned (Figure 3). Corresponding amino acid residues, R88 and D97, were also found in studied halophyte protein SaNAR2.2 (Fig. 3).

According to protein topology prediction, only one C-terminal transmembrane segment is present in SaNAR2.2, while central part of SaNAR2.2 with conservative amino acid residues important for the interaction of the two partner proteins is located in the extracellular space (Figure 2b). Therefore, it is reasonable to assume that interaction of SaNAR2 and SaNRT2.1/SaNRT2.5 occurs in the extracellular space. This assumption is confirmed by in silico model of interaction between SaNAR2.2 and SaNRT2.5, which was created using AlfaFold3 software (Figure 4с). Besides, the 3D-model of SaNAR2.2/SaNRT2.5 heterodimer suggests that transmembrane C-terminal region of SaNAR2.2 protein may also be involved in this interaction.

In studies of *SaNAR2.2* expression under various ionic environmental conditions, results were obtained that suggest the involvement of SaNAR2.2 protein in high-affinity nitrate uptake by *S. altissima* plants. Apparently, the role of SaNAR2.2 protein as a participant of high-affinity nitrate uptake system was particularly significant under saline conditions. *SaNAR2.2* was expressed nearly exclusively in *S. altissima* roots (Figure 6a), similarly to *NAR2* genes from other plant species [23, 24, 27–30, 31]. In the absence of salt in the nutrient solution the expression levels of *SaNAR2.2* gene at low (0.5 mM) and high (15 mM) nitrate concentrations were similar (Figure 6a). However, under increase of external NaCl concentration up to 500 mM, the expression of SaNAR2.2 was slightly decreasing at high nitrate concentration in the medium but significantly increased at low nitrate concentration (Figure 6b). With further increase in salt concentration in the nutrient solution up to 750 mM, the expression of *SaNAR2.2* decreased at low nitrate. Under low nitrate, the expression of *SaNRT2.1* and *SaNRT2.5*, which encode high-affinity nitrate transporters of the NRT2 family in *S. altissima* and also expressed in *S. altissima* roots [48], changed in a similar way depending on changes in external NaCl concentrations (Figure 6 c, d). It was demonstrated earlier [48] that at low external nitrate concentrations the growth of *S. altissima* plants was predictably reduced compared to plants grown at high concentrations of nitrate; at the same time, plant weight and nitrate accumulation in the aboveground organs of *S. altissima* were maximal at a high salt concentration of 500 mM NaCl (further increase in NaCl concentration in the medium to 750 mM led to reduced biomass and lower nitrate accumulation in the aboveground organs). This indicates the effective functioning of high-affinity nitrate uptake systems under conditions of strong competition between nitrate and chloride for nitrate-transporting mechanisms. Similar expression patterns of the three genes, *SaNRT2.1*, *SaNRT2.5*, and *SaNAR2.2*, depending on the changes in nitrate and salt concentrations in the environment, were demonstrated. Obtained data allowed us to suggest that there is a functional relationship between the products of these genes as has been shown for orthologous proteins from other plant species [22–26].

It is worth to note that under low nitrate and high salinity the expression of *SaNAR2.2* reached an extremely high level, which was almost twice more than expression of the house-keeping gene *SaeEF1ɑlfa*, while the expression levels of *SaNRT2.1* and *SaNRT2.5* were significantly lower (Figure 6). The expression of *SaNRT2.5* was almost 20 times lower than the expression of *SaNAR2.2*, and the expression of *SaNRT2.1*, in turn, was 10 times lower than the expression of *SaNRT2.5*. Such differences in the expression levels of the *SaNRT2.1*/*SaNRT2.5* genes and the *SaNAR2.2* gene, the product of which forms a complex with high-affinity nitrate transporters of the SaNRT2 family, may probably be explained by the fact that NAR2s play multiple roles in addition to being a component of the high-affinity nitrate transport system in plants [27]. For example, it has been shown that both high- and low-affinity nitrate transport were greatly impaired by *OsNAR2.1* knockdown in rice plants [29]. This suggests the involvement of NAR2 proteins in low-affinity nitrate transport, but the mechanisms of such involvement have not yet been investigated. NAR2 proteins are also involved in the root growth regulation. Knockdown of *OsNAR2.2* significantly repressed the primary root elongation and severely reduced the number of lateral roots under low nitrate concentrations [42, 43]. These findings allowed the authors to suggest that NAR2 affects the root growth and development of rice; the effects are most likely achieved through a combination of roles in both NO^3̶^ and auxin signaling. We can assume that in *S. altissima*, the role of SaNAR2.2 is not limited to participation in high-affinity nitrate transport as well.

It can also be assumed that the significantly higher *SaNAR2.2* expression in comparison to the expression of the nitrate transporter genes *SaNRT2.1* and *SaNRT2.5* is, in addition to other factors, related to the stoichiometry of formation of the heterologous complex of SaNRT2s and SaNAR2.2 proteins in *S. altissima*. For *A. thaliana*, it has been shown that a 150kDa plasma membrane complex of AtNRT2.5 and AtNAR2.1 is the major contributor to high-affinity nitrate influx [73]. Also, the AtNRT2.1 forms the 150 kDa oligomeric complex with the AtNAR2.1 protein [25]. The 150kDa oligomer has been proposed to contain a tetramer consisting of two subunits each of AtNRT2 and AtNAR2.1 [27]. It is possible that *S. altissima* has a different stoichiometry for the formation of the heterologous complex of proteins SaNAR2.2 and SaNRT2, requiring more than one molecule of SaNAR2.2 per molecule of SaNRT2. If so, then it could be one of the reasons of incomplete functional complementation of the yeast mutant strain *Δynt1* by SaNRT2.1/SaNRT2.5 and SaNAR2.2 heterologous expression (Figure 7). In yeast, the expression level of *SaNAR2.2* could be insufficient for the formation of a functional nitrate transport complex. In our work we used integrative yeast plasmids, the number of copies of which integrated into the yeast genome is generally random and low. It is also possible that, at a sufficient level of expression, only a small fraction of the synthesized heterologous proteins reached the plasma membrane of yeast cells. Indeed, we observed that heterologous *Suaeda* proteins synthesized in the yeast cells remained localized in the internal space of the yeast cell and did not reach the plasma membrane (Figure 8). These observations may indicate that *Suaeda* proteins require post-translational modifications to exhibit their transport activity that occur in the native system but are absent in yeast cells. For example, it has been shown that phosphorylation of serine residues in the N-terminal region of the AtNRT2.1 proteins in *A. thaliana* was necessary to enhance the stability of the AtNRT2.1-AtNAR2.1 protein complexes and increased its transport activity [74, 75].

## Conclusions

The coding sequence of *NRT3/NAR2* family gene, *SaNAR2.2,* was cloned from the euhalophyte *Suaeda altissima.* The terrestrial euhalophyte is unique from the point of its extremely high salt tolerance: it is able to grow and proliferate at salinities up to 1 M NaCl. Under high salinity the euhalophyte efficiently takes up nitrate under low nitrate concentrations in the medium at the expenses of high affinity nitrate uptake system. The system is formed by NRT2 family proteins while gene *SaNAR2.2* encodes partner protein SaNAR2.2 for the functioning of NRT2 family proteins. In silico analysis and bimolecular fluorescence complementation (BiFC) assay in leaves of *Nicotiana benthamiana* evidence for interactions of SaNAR2.2 with high affinity nitrate transporters of the euhalophyte SaNRT2.1 and SaNRT2.5. Expression analysis indicated that *SaNAR2.2* is expressed nearly exclusively in roots. Moreover, at low (0.5 mM) nitrate in the nutrient medium the expression of *SaNAR2.2* increased about ten times at 500 mM NaCl in the nutrient solution; this increase coincided with the increase in expression of *SaNRT2.1* and *SaNRT2.5*. Opositedly, expression of these three genes decreased at high (15 mM) nitrate and salinity. The results obtained point to an important role of SaNAR2.2 in nitrogen uptake under salinity in the halophyte.

## Author Contributions

P.L.G., K.D.E.: conceptualization, validation, writing - original draft preparation, data curation; K.D.E., K.A.O., N.O.I., R.E.I. and R.A.V.: methodology, investigation, formal analysis; P. L.G., K.D.E. and V.V.S.: formal analysis, writing - review and editing. P.L.G.: supervision; P.L.G.: funding acquisition. All authors have read and agreed to the published version of the manuscript.

## Institutional Review Board Statement

Not applicable.

## Informed Consent Statement

Not applicable.

## Data Availability Statement

All data included in this study are available upon request by contact with the corresponding authors. The seeds of *Suaeda altissima* are available from the authors on request. The cloned *SaNAR2.2* coding sequence was deposited in GenBank (*Suaeda altissima* nitrate assimilation related protein mRNA, ID: OR828749.1).

## Acknowledgments

We are grateful to Mikhail O. Agafonov (A.N. Bach Institute of Biochemistry of Federal Research Center“Fundamentals of biotechnology”, Russian Academy of Sciences, Moscow, Russia) for kindly provided pCCUR2 and pCHLX plasmids and to Sergey N. Lomin (K.A. Timiryazev Institute of Plant Physiology, Russian Academy of Sciences, Moscow, Russia) for kindly provided pSPYNE-35S and pSPYCE- 35S plasmids. The authors sincerely thank Svetlana Bagirova for critical reading and comments on the manuscript.

