## Supplementary tables 1-2 for "Molecular Cloning, *In Silico* Analysis and Expression of plasma membrane-associated NAR2 Protein, SaNAR2.2, from Euhalophyte *Suaeda altissima*"

| **Primer name** | **Sequence** |
| --- | --- |
| ***qRT-PCR*** | |
| SaNRT2.1_F1 | GTGCTTCTGTATGGATACTCT |
| SaNRT2.1_R1 | ATTCCAGCCGTGTGTAG |
| SaNRT2.5_F2 | TACTGGATGAGCTCCATGTT |
| SaNRT2.5_R2 | ACATTGTAGATTAGAGGCATCAC |
| SaNRT3.2_F1 | CATGGAGGAGAACCGAAG |
| SaNRT3.2_R1 | CTTGGTAGGTTACTGTGTCATC |
| SaeEF1alfa_F1 | TGAGATGTGTGGCAATCC |
| SaeEF1alfa_R1 | GTTGCTTCTGACTCCAAGAAT |
| ***5’ RACE (I round)*** | |
| SaNRT3.2_R | ACAATTAAGAAGCCCTTTTAGCCT |
| ***3’ RACE (I round)*** | |
| SaNRT3.2_F | AAAAGTTAAGTAGTAATGGCGATTCGT |
| ***5’ RACE (II round)*** | |
| SaNRT3.2_R1 | CTTGGTAGGTTACTGTGTCATC |
| ***3’ RACE (II round)*** | |
| SaNRT3.2_F1 | CATGGAGGAGAACCGAAG |
| ***Full-length SaNAR2.2 CDS amplification*** | |
| SaNRT3.2_F3 | ATGGCGGTTCGTGGATTAATC |
| SaNRT3.2_R3 | TTAAGAAGCCCTCTTAGCCTTACTC |
| ***pYNR1 amplification*** | |
| pCCUR2-pYNR1-F | CTGCTTTTGTCTTGTGGTACTATGGGGCTCCATATATCGTATGAC |
| pYNR1-SaNRT3.2_R | TCCACGAACCGCCATAGATCTTACAACTATCCAAAGTTCGTGAGG |
| ***tYNR1 amplification*** | |
| tYNR1-SaNRT3.2_F | GCTAAGAGGGCTTCTTAAGCTAGCCCTAAGCGAAATCGAAATCAAAACT |
| pCCUR2-tYNR1-R | CTAAAGGGAACAAAAGCTGGAATTCAGTATTTCAGAATCATGACCC |
| ***GFP amplification*** | |
| GFP_Gib_F | CCAGCGGCCGCCATGAGCAAGGGCGAGG |
| tYNR1_GFP_R | CGCTTAGGGCTAGCTTACTTGTACAGCTCGTCCATG |
| ***mCherry amplification*** | |
| mCherry_Gib_F | CCAGCGGCCGCCGTGAGCAAGGGCGAG |
| tYNR1_mCherry_R | TCGCTTAGGGCTAGCTTACTTGTACAGCTCGTCC |
| ***pCHLX-SaNRT2.5 linearization*** | |
| SaNRT2.5_Gib_R | CATGGCGGCCGCTGGGACACGATTAACTGTACTGCC |
| tYNR1_F20 | GCTAGCCCTAAGCGAAATCG |
| ***pCCUR2-SaNAR2.2 linearization*** | |
| mCherry_SaNRT3.2_R | CACGGCGGCCGCTGGAGAAGCCCTCTTAGCCTTAC |
| tYNR1_F20 | GCTAGCCCTAAGCGAAATCG |

**Table S1.** List of the primers used in the research.

**Table S2**. **List of NAR2 proteins from the phylogenetic tree of Figure 2.**

| **Plant species** | **Protein name** | **GenBank ID** |
| --- | --- | --- |
| *Amaranthus tricolor* | **AtrNAR2.2** | XP_057534669.1 |
| *Arabidopsis lyrata* | **AlNAR2.1** | XP_020871335.1 |
| *Arabidopsis thaliana* | **AtNAR2.1** | NP_199831.1 |
| *Arabidopsis thaliana* | **AtNAR2.2** | NP_001190826.1 |
| *Asparagus officinalis* | **AoNAR2.1** | XP_020246811.1 |
| *Beta vulgaris* | **BvNAR2.2** | XP_010667500.1 |
| *Brassica rapa* | **BrNAR2.1** | XP_009134048.2 |
| *Capsicum chinense* | **CchNAR2.2** | PHU23681.1 |
| *Chenopodium quinoa* | **CqNAR2.2** | XP_021720161.1 |
| *Eutrema salsugineum* | **EsNAR2.1** | XP_006402168.1 |
| *Eutrema salsugineum* | **EsNAR2.2** | XP_024005789.1 |
| *Glycine max* | **GmNAR2.1** | XP_003549822.1 |
| *Hordeum vulgare* | **HvNAR2.1** | AAP31850.1 |
| *Hordeum vulgare* | **HvNAR2.2** | AAP31851.1 |
| *Iris pallida* | **IpNAR2.1** | KAJ6792152.1 |
| *Medicago truncatula* | **MtNAR2.1** | XP_013457733.1 |
| *Oryza sativa* | **OsNAR2.1** | Q6ZI50.1 |
| *Oryza sativa* | **OsNAR2.2** | Q7XK12.1 |
| *Perilla frutescens* | **PfNAR2** | KAH6794745.1 |
| *Phtheirospermum japonicum* | **PjNAR2.1** | GFQ07351.1 |
| *Salvia hispanica* | **ShNAR2.1** | XP_047977271.1 |
| *Sesamum indicum* | **SiNAR2.1** | XP_020548343.1 |
| *Solanum pennellii* | **SpNAR2.2** | XP_015069077.1 |
| *Spinacia oleracea* | **SoNAR2.2** | XP_021863523.1 |
| *Suaeda altissima* | **SaNAR2.2** | WPH61291.1 |
| *Suaeda fruticose* | **SfNAR2.2** | Transctiptome assembl. |
| *Suaeda glauca* | **SgNAR2.2** | Transcriptome assembl. |
| *Triticum aestivum* | **TaNAR2.2** | AAV35211.1 |
| *Triticum aestivum* | **TaNAR2.1** | AAV35210.1 |
| *Vigna radiata* | **VrNAR2.1** | XP_014490580.1 |
| *Zostera marina* | **ZosmaNAR2.1** | KMZ59910.1 |

>AtrNAR2.2

MAVRGLIIFAMIFSVMVTTCCGKVMFSHLANTLTVSTSPKGKVDLKAGVDSITITWGLNKNGSKVDTKSY

KNIEVKLCYSKESQKDRPWRKTEEELDRDKTCQFSVVKKEYNVTTDSFTYKVKKDVPTAHYFVRVYVRDG

PDGKQIAYGQTNGVDLNVTGISGRSSSIDIAASVFSTFSVVSLAFFFYLEKKKARKEEAK

>AlNAR2.1

MAIHKILFASLLICSLIQSSHGAEKVRLFKELDKGALDVTTQPSRQGDGVVLDAGKDTLNITWKLSAIGS

KREAEFKIIKVKLCYAPPSQLDRPWRKTHDELFKDKTCPHKIIAKPYDKTPQSTVWTVERDIPTGTYFVR

AYAVDAIGHEVAYGQSTDDAKTTNLFSVQAISGRHTSLDIASICFSVFSVVALVVFFVNEKRKAKIEQSK

>AtNAR2.1

MAIQKILFASLLICSLIQSIHGAEKVRLFKELDKGALDVTTKPSREGPGVVLDAGKDTLNITWTLSSIGS

KREAEFKIIKVKLCYAPPSQVDRPWRKTHDELFKDKTCPHKIIAKPYDKTLQSTTWTLERDIPTGTYFVR

AYAVDAIGHEVAYGQSTDDAKKTNLFSVQAISGRHASLDIASICFSVFSVVALVVFFVNEKRKAKIEQSK

>AtNAR2.2

MRHMCDTHLVVLPYQDHILIRIMAIHTLLFVSLLIFSLIESSSGGKKDRLFTDLQNSIEVTAKPVKDSGV

LEAGKDMVTITWKLKSSSAKVDTDTAFKTIQVKLCYAPISQVDRPWRKTDNKLFKDRSCPHEIVSKAYDK

TPQSLDWTIGLDIPTGTYFVRAYGIDGDGHEVAYGQSTDEGRTTNLFSVHAISGHHVGLDIASTFFSVFS

VVSLFVFFVMEKRKAKLEQRE

>AoNAR2.1

MENQRRNNQEMAASSLFFSPATILGLLLLFSIGSSVEGVLFSKLKQTLIVTASPNHGQVLKAGEDKLTVS

WALNSTVSDSSYKQVKVILCYAPVSQKDRGWRKTVDDLSIDKTCQFDIITKPYTKSKTTFEYMVKRNVPT

ATYFIRAYVLDFSENEVAYGQTTDAAKTTKFFDIIGISGRTASLDIAAGCFSAFSILSLVYFFVKEKRMA

KK

>BvNAR2.2

MGVRGLIIFAAVLCSLVATCYSKGVFADLRNTLTVSASPSGRVDLRAGIDQITVSWRLNRNISNTDSATY

SKVDVKLCYHLESQKDRPWRRTEEDLSRDKTCQFSMVKKDYNSSTDSVTYTVKKNIPTAHYFVRVYVRNA

DNREIAYGQTNGLDLNIKGISGRSTSIDVAASVFSGFSVLSLAFFFFLEKRKAKKLTS

>BrNAR2.1

MCTPPFISNIKSQQTKAYSSSLSKYSDSRISMAIHKSLFASLLICLLFQISHGATKERLFSDLEKGALEV

TAKPSREGVLDAGIDKLSITWKLSSTATKEAEFTTIKVKLCYAPVSQVDRPWRKTENELFKDKSCPHKII

TRAYDKSPQSFEYTLERDIPTGTYFVRAYAVDAKDHEVAFGQSTNEAKSTNLFSVQAISGRHKSLDIASV

CFSVFSVLALLVFFVNEKRKAKIEQSK

>CchNAR2.2

MASTTAIFLATLVICCSLSSSHAEILFSSLKKSLEVTVKHRAGVLMAGEDTLEIDWFLNKTFPAGTDSAY

KTIKLKLCYAPISQKDRAWRKTEDHLKKDKTCLFEIDSTPYKSSNNKFNWTIERDVPTGTFFVRAYILNG

DGHEIGYGQNTDDKKVNNLFDIQAISGRHATLDICSVVFSVFAVVSLFGFFYMEKRNAKASK

>CqNAR2.2

MAVRGLIIFSVLLSSLVASCYGTGLFKDLSNTLTVSTTPTGKVNLKAGKDQITVTWGLNRNVSKLDTSAY

KQVEVKLCFLAESQVDRPWRKTEDELARDKTCQFLVVKKDFTSSSDSFTYTVKKDVPTAHYFVRVYVRDG

PDGKQLAYGQTTGLDLSVKSISGRSASIDIAASIFSAFSVLSLAFFFYLEKKKARRAT

>EsNAR2.1

MAIHKILFASLFICSLIQSSHGATKERLFSDLEKGAFEVTTKPSREGEGVVLDAGIDKLSITWKLNSTAK

EAEFKTIKVKLCYAPISQVDRPWRKTENELFKDKSCPHKIVARPYDSSVKTAQTIDYTLERDIPTGTYFV

RAYAVDAHDHEVAFGQSTDEDKKTNLFSVQAISGRHKSLDIASICFSVFSVLALLVFFVNEKRKAKIEQS

K

>EsNAR2.2

MAIHSFLFISLLILSSLESSSGTTKDRLFFTNLQNTLEVTAKPIRDGVLESGKDIIRVTWKLKSSVKIDV

DAAIKTVDVRLCYAPISQVDRPWRQSHNELFKDKTCPYKIVSKPYDKISQSLNWIIERDIPTGTYFVRVY

GIDANGHEVAYGQXSNAEKTTNLFSVEAIDGRHISLDAASICFSIFSVVALLVFFMRERRRNPI

>GmNAR2.1

MAAQGPVVVASLLIFCLAGSCYGKVHFSSLKRTLDVTASPKQEQVLEAGLDKITVTWALNKTLPAGTDSA

YKTIKLKLCYAPISQKDRAWRKTEDELKRDKTCQHKIVAKPYDASNKTVQRFEWLVERDVPKATYFVRAY

AFDSNDEEVAYGQTTDAKKSTNLFEINAVSGRHASLDICSICFSAFSVVSLFVFFYIEKRKGKASSSK

>HvNAR2.1

MARSELVMVLLVVVLAAGCCTSAGAVAYLSKLPVTLDVTASPSPGQVLHAGEDVITVTWALNTTQAGKDA

DYKNVKVSLCYAPVSQKEREWRKTHDDLKKDKTCQFKVTQQAYPGAGKVEYRVALDIPTATYYVRAYALD

ASGTQVAYGQTAPTAAFNVVSITGVTTSIKVAAGVFSAFSVASLAFFFFIEKRKKNN

>HvNAR2.2

MARSDLVMALLVAVLAAGCCASAGAVAYLSKLTVTLDVTASPTPGQVLHAGEDVITVTWALNATRPAGDD

AAYKSVKVSLCYAPASQKEREWRKTHDDLKKDKTCQFKVTQQPYAAGAAGGRVEYRVALDIPTAAYYVRA

YALDASGTQVAYGQTAPATAFDVVSITGVTTSIKVAAGVFSTFSVVSLAFFFFIEKRKKNN

>IpNAR2.1

MGDLFSIPVLFFLLFLLVGTAAGEGVLLSTLPRTLIVTASPKQGQVLKAGEDHITVSWGLNKSLSASTDD

AYKNVKVMLCYAPVSQTDRGWRKTNDDLKKDKTCQFEVTVQPYAKSNSSFEYKIKRDVPSATYFLRAYLL

DSSDTKVAYGQTTDAHKSTNLFEIVGISGRSASLDVAAGCFSAFSVVSLLFFFVIEKRKAKKSLN

>MtNAR2.1

MAAHKLVVASLLLCCLAEICYGKDLFSSLKRTIDVTASPKQGQVLLSGVDKISGTWALNKTFPAGTDSSY

KTIKLKLCYAPISQKDRAWRKTEDELSRDKTCQHKMLAMPYNASNKTVQTFEWLIQRDVPQATYFVRAYA

FDSNDKEVAYGQTTNAGKSTNLFEINAISGRHATLDICSVVFSAFSVVSLGVFFYIEKRKGKSPKQ

>OsNAR2.1

MARLAGVAALSLVLVLLGAGVPRPAAAAAAKTQVFLSKLPKALVVGVSPKHGEVVHAGENTVTVTWSLNT

SEPAGADAAFKSVKVKLCYAPASRTDRGWRKASDDLHKDKACQFKVTVQPYAAGAGRFDYVVARDIPTAS

YFVRAYAVDASGTEVAYGQSSPDAAFDVAGITGIHASLKVAAGVFSTFSIAALAFFFVVEKRKKDK

>OsNAR2.2

MARFGAVIHRVFLPLLLLLVVLGACHVTPAAAAAGARLSALAKALVVEASPRAGQVLHAGEDAITVTWSL

NATAAAAAAGADAGYKAVKVTLCYAPASQVGRGWRKAHDDLSKDKACQFKIAQQPYDGAGKFEYTVARDV

PTASYYVRAYALDASGARVAYGETAPSASFAVAGITGVTASIEVAAGVLSAFSVAALAVFLVLENKKKNK

>PfNAR2

MAIQGFLVASILISCFVATCYGVTFSSLQNTLVVTSSPHSDQELKAGEDKITVTWSLNTTFPSGTDSTYK

MVKVKLCYAPISQKDRGWRKTVDNLKKDKTCQHKIVAKPYTASNNTFTYTVQRDVPTATYFVRAYVVNAA

DEEVAFGQSTDSHKATNLFKIQAISGRHISLDIASVCFSAFSIVSLFGFFYMEKRKAK

>PjNAR2.1

MANHVSVIALISLSCLVATCYGNVLFSTLNPTLIVTASPSPPGQVLKAGEDTITVTWSYNSSFPSSTDSD

YRTVKIMLCYAPESQQDRAWRKTVDNLKKDKTCQHKIVSRPYSRSGNNTFTYTVQRDIPTATYFVRAYAL

NAHDTQLAYGQTTNAQKTTNLFGIQAITGRHVSLDIASICFSGFSILSLFGFFYMEKRKAKSQSK

>ShNAR2.1

MAINRVIVATILLSCLAATCYAITFSSLQRTLDLDSSVQAGQVLKAGEDKITVSWKVNATYPAGTESSYK

TVKVLLCYAPVSQKDRGWRKTVDLLKKDKTCQHTIVERAYSASDNSVTYTVKRDVPTATYFIRAYVYNAA

GEEVGYGQTTNADKSANLFAVEAVTGRHVSLDIASACFSAFSVVSLFGFFFMEKRKAKASN

>SiNAR2.1

MAIQGSIVASILLSYLVAACYGATFSSLQNTLVVTASPQAGQVLKGGEDNITVTWSLNNTFPAGTDSDYK

TVKIKLCYAPISQQDRAWRKTEEELKRDKTCQHKIVAKPYSPSTNTFTWTVQRDVPTATYFIRAYVHNSA

DKEVAFGQTTDSHKATNLFEVQAISGRHVSLDIASVCFSAFSVVALFGFFFLEKRKAKASQKS

>SpNAR2.2

MAISTRAIFVATIVICSFLAVCEGEILFSTLKKSLEVKADHKAGVLMAGEDALTLKWSLNNTLPAGSDSG

YKSVKLELCYAPISQKDRAWRKTEDHLKKDKTCQFKIATVPYKSSNNNYNWTIKRDIPTATYFVRAYVLN

GNGEEIGYGQNTDDKKVDNLFSIQAISGRHATLDICSVVFSAFSVVSLFAFFYMEKRKARAGK

>SoNAR2.2

MAVRGLIIFVALLSSLAASCYGVMFHELAHSLTVSSTPSGQANVKAGKDQITVTWALNRSISGVDTSIYR

EVEAKLCYHLESQKDRPWRKTEEEMARDKTCQFAIVKRPYTSSSDSVTYTIKKDVPTAHYFIRVYVRDGP

GGKLIAYGQTTGLDLFVAGISGRSASIDIAASIFSAFSVISLGFFFYLEKKKSKRAT

>SaNAR2.2

MAVRGLIIFVALLSSFVVACYGEGLFSELKQTLTVSASPTGKVSLIAGKDEITVTWGLNSTSTKTTNYKN

VEVKLCYLEESQKDRPWRRTEDDLKRDKTCQFLVAEKEFTTKDDTVTYQVKKDVPSAHYFIRAYVTDGPK

GKLIAYGQTTGLDLKVKGISGRSLAIDVAASVFSGFSVLSLVFFYFMEKSKAKRAS

>SfNAR2.2 MAVRGLIIFVALLSSFVVACYGEGLFSELKQTLKVSSSPTGKVFLMAGKDEITVTWGLNSTSTKTTNYKNVEVKLCYLEESQKDRPWRKTEDDLKRDKTCQFLVAEKEFTTKDDTVTYKVKKDVPSAHYFIRAYVSDGPKGKLIAYGQTTGLDLQVKGISGRSLAIDVAASVFSGFSVLSLVYFYFMEKSKAKRAS

>SgNAR2.2 MAIRGLIIFGALLSSFVVACYGEGLFSDLKQTLSVSASPTGKVFLTAGKDEITVTWGLNSTSTKTTNYKNVDVKLCYLAESQKDRPWRRTEDDLKRDKTCQFLVAEKEFNTKGDKVTYKVKKDVPSAHYFIRAYVKDGPKGKLIAYGQTTGLDIEVNGISGRSLAIDVAASVFSGFSVLSLVFFYFMEKRKAKRAS

>TaNAR2.2

MAQSKLVMALLVAVLAAGCCASAGAVAYLSKLPVTLDVIASPSPGQVLHAGEDVITVTWALNASRPAGDD

AAYKNVKVSLCYAPASQKEREWRKTHDDLKKDKTCQFKVAQQPYAGAGGRVEYRVALDIPTATYYVRAYA

LDASGTQVAYGQTAPAAAFNVVSITGVTTSIKVAAGVFSTFSVVSLAFFFFIEKRKKNN

>TaNAR2.1

MARQGMVTALLLVVLAAGCCASAGAVAYLSKLPVTLDVTASPSPGQVLHAGEDVITVTWALNASQPAGKD

VDYKNVKVSLCYAPVSQKEREWRKTHDDLKKDKTCQFKVTQQAYPGTGKVEYRVALDIPTATYYVRAYAL

DASGTQVAYGQTAPSSAFNVVSITGVTTSIKVAAGVFSAFSVASLAFFFFIEKRKKNN

>VrNAR2.1

MAAHRLVVVSLLIFCFAGSCYGKVHFSSLKKTLDVTASPKQGQVLEAGTDKITVTWALNNTLPTGTDSAY

KTIKVKLCYAPISQKDRAWRKTEDELSRDKTCQHKIVAKPYDASNRTVQRFEWVIERDVPKATYFVRAYA

FDSNGEEVAYGQTTDAKKSSNLFEINSISGRHASLDICSVCFSAFSVVALFVFFYIEKRKAKASSSSK

> ZosmaNAR2.1

MYSSFLSVAVSVAVFAVVCCVSPLVDAEVMFSSLPENLIVDASPIQGQVLKAGEDNITVSWMLKTNLAPG

VDAKYQSIKIKLCYGPSSQVDRGWRKTRDNLKKDKTCQFNVGESLIYNSNAAQMEQRVTYTISREVPTAM

FFVRAYVLDSNGKKIAYGQSTNAEKNTNLFEVEGITGRNTSLNIAATCFSAFSLMSMTGFYIADRRRKNK
